# Implantation-Competent Blastocyst-Like Structures from Mouse Pluripotent Stem Cells

**DOI:** 10.1101/309542

**Authors:** Cody Kime, Hiroshi Kiyonari, Satoshi Ohtsuka, Eiko Kohbayashi, Michio Asahi, Shinya Yamanaka, Masayo Takahashi, Kiichiro Tomoda

**Affiliations:** Gladstone Institute of Cardiovascular Disease, San Francisco, CA 94158, USA; Lab of Retinal Regeneration, RIKEN Center for Biosystems Dynamics Research, Kobe 650-0047, Japan; Animal Resource Development Unit and Genetic Engineering Team, RIKEN Center for Life Science Technologies, Kobe 650-0047, Japan; Department of Life Science, Medical Research Institute, Kanazawa Medical University, Ishikawa 9200293, Japan; Second Department of Internal Medicine, Osaka Medical College, Osaka 569-8686, Japan; Department of Pharmacology, Faculty of Medicine, Osaka Medical College, Osaka 569-8686, Japan; Center for iPS Cell Research and Application (CiRA), Kyoto University, Kyoto 606-8507, Japan; Lead Contact

**Author notes:** **Corresponding Authors**: Correspondence should be addressed to C.K. or K.T.

**Keywords:** pluripotent stem cells, early embryo, implantation, cell biology, totipotency, reproduction, cell plasticity, blastocyst

## Abstract

Soon after fertilization, the few totipotent cells of mammalian embryos diverge to form a structure called the blastocyst (BC). Although numerous types of cells, including germ cells and extended pluripotency stem cells, have been generated from pluripotent stem cells (PSCs) *in-vitro*, generating functional BCs only from PSCs has not yet been reported. Here we describe induced self-organizing 3D BC-like structures (iBCs) generated from mouse PSC culture *in-vitro*. Resembling natural BCs, iBCs have a blastocoel-like cavity and were formed with outer cells that are positive for trophectoderm lineage markers and with inner cells that are positive for pluripotency markers. iBCs transplanted to pseudopregnant mice uteruses implanted, induced decidualization, and exhibited growth and development before resorption, demonstrating that iBCs are implantation-competent. iBC production required the transcription factor Prdm14 and iBC precursor intermediates concomitantly activate the MERVL totipotency related cleavage stage reporter. Thus, our system may contribute to understanding molecular mechanisms underpinning totipotency, embryogenesis, and implantation.

**HIGHLIGHTS:**

-Pluripotent cells self-organize blastocyst-like structures in defined conditions.
-Structures have several extraembryonic and embryonic characteristics of blastocysts.
-Structures can implant in the uterus and grow before resorption.
-Totipotency is implicated concomitantly at loci that originate induced blastocysts.

## INTRODUCTION

During early mammalian development, a fertilized egg (zygote) completely intersects the animal life cycle upon zygotic genome activation (ZGA): the event where gamete totipotent genomes of the pronucleus are epigenetically activated and rapidly enter cleavage (Seydoux and Braun, 2006; Wu et al., 2017). The zygote cleaves and symmetry later bifurcates to form the blastocyst (BC) in preparation for implantation and differentiation. The BC is a 3D ball-like structure of three characteristic parts: the extraembryonic (ExEm) outer layer of trophectoderm (TE) lineage cells, the pluripotent cells of the inner cell mass (ICM), and a fluid filled cavity called the blastocoel. Upon implantation, trophoblasts contribute to the ExEm tissue of the placenta, and the ICM gives rise to embryo proper and some ExEm tissues. Emerging trophoblasts and pluripotent cells result from the first differentiation event in mammalian development, initiated in cleaving totipotent cells that begin to polarize just before the BC forms (Hirate et al., 2015; Nishioka et al., 2009; Yu et al., 2016; Stephenson et al., 2010).

Implantation is crucial to natural development and establishes the physical connection between the mother and early embryo that supports embryonic (Em) development through the rest of the pregnancy. Implantation is tightly regulated at several molecular and cellular levels: apposition, adhesion and invasion of the TE lineage cells, and subsequent decidualization of the endometrial wall of the uterus. ExEm tissues thereafter increase growth and differentiation while the ICM differentiates to form the embryo proper and additional ExEm tissues. Implantation-competent BCs require TE cells expressing Cdx2 (Meissner and Jaenisch, 2006), and molecular mechanisms involving Lpar3, Lif, Bmp, and others signal the interface between the TE and the receptive uterus (Cha et al., 2012; Wang and Dey, 2006). These events must occur in a short developmental window: failed implantation is a major cause of early pregnancy loss in humans (Norwitz et al., 2001; Cha et al., 2012). Defective embryos also fail later and begin the resorption process in which maternal immune cells degrade the embryo (Cossée et al., 2000; Flores et al., 2014).

The zygote and cleavage stages exhibit true totipotency, isogenically preceding all ExEm (vegetal) and Em (animal) cell bi-directional development toward entire organisms. From plant tissue cultures, specific cytokine, vitamin, and plant hormone (auxins) ratios are adjusted to induce totipotent transient cells for propagating isogenic embryos (Steward et al., 1958). In mammals, isogenic 3D BCs from differentiated cells are both attractive and elusive. Recent progress in PSC research challenges this barrier by generating functional germ cells that give rise to offspring by *in-vitro* fertilization (Hikabe et al., 2016), and some reports show stem cells with extended/bi-directional pluripotency in chimeric mice (Macfarlan et al., 2012; Yang et al., 2017). However, experiments inducing implantation-competent isogenic BCs entirely from PSCs are unprecedented.

In reprogramming and conversion experiments with specific cytokines, nutrient, and lipid (Kime et al., 2016), we frequently observed tissues and hemispheres that resemble BCs. Inspired by these observations, we developed a stepwise regime to readily induce BC-like structures from PSCs *in-vitro*, which we term iBCs. iBCs demonstrate implantation-competence since transplant into pseudopregnant mice induced focal decidualization in the uterus, recruited a maternal blood supply, and expanded the embryonic cavity. Some implanted iBCs produced many cell types similar to implanted embryos but failed to develop further due to embryonic resorption. A live pluripotency reporter suggests pluripotency is partially recovered in the putative ICM of iBCs, and some iBC express *Zscan4*. Utilizing the murine endogenous retrovirus (*MERVL*) live totipotency-related reporter, we found iBC precursors and cells where iBCs originate may indicate ZGA mechanisms (Macfarlan et al., 2012; Wu et al., 2017). Further analysis of Yap protein distribution in iBC precursors and early iBCs was similar with cleavage stage cells polarizing through compaction toward emerging blastocysts (Nishioka et al., 2009; Stephenson et al., 2010; Bedzhov et al., 2014). We anticipate this approach may lead to simplified isogenic embryo production for research, medicine, and uncovering the intricacies of totipotency and implantation.

## RESULTS

### Efficient Conversion to Naive PSCs Produces Blastocyst-like Hemispheres

*In-vitro* pluripotency is characterized in two distinct states: a post-implantation epiblast state (primed) and a pre-implantation blastocyst ICM state (naive). Primed female PSCs have one active and one inactive X chromosome (Xa/Xi), but naive female PSCs have two Xas (Xa/Xa) (Payer et al., 2011). We previously presented defined conditions that enhanced iPS cell reprogramming, and in primed to naive PSC conversion experiments, those conditions formed structures resembling early embryonic material. In the naive conversion experiments, we used a primed female mouse epiblast stem cell (mEpiSC) line that harbors a silent green fluorescent protein (GFP) transgene on the Xi chromosome (Xi-GFP, XGFP-). GFP is expressed upon reactivation of Xi to Xa (Xa-GFP, XGFP+), a hallmark of the ICM and often of cleavage stage cells (Kime et al., 2016; Monk and Harper, 1979; Okamoto et al., 2004; Bao et al., 2009).

Robust naive conversion frequently produced hemispheres with morphological characteristics we suspected to resemble BCs (Figure 1). XGFP+ cell clusters, which we found to be naive cells (Kime et al., 2016), were polar and internal to fluid-filled hemispherical domes of distinct large flat cells with ectodermal morphology resembling trophoblasts. Using immunocytochemistry, we found that these hemispheres initiated the fluid-filled cavity and had NANOG+XGFP+ inner cells with no bright DNA-stain punctae, which may indicate loss of heterochromatin usually found in scarce transient Zscan4+ 2C-like state cells in mouse naive PSCs (Akiyama et al., 2015; Wu et al., 2016). The inner cells were surrounded by spheroid DNA-stained punctae-enriched NANOG+XGFP− outer cells. These cell distributions and expression characteristics were similar to early differentiating cells of the morula. Also, the most-outward cells were flattened and NANOG-XGFP−, characteristic of lineage committed TE. (Figure 1A, Figure S1A).

**FIGURE 1:**
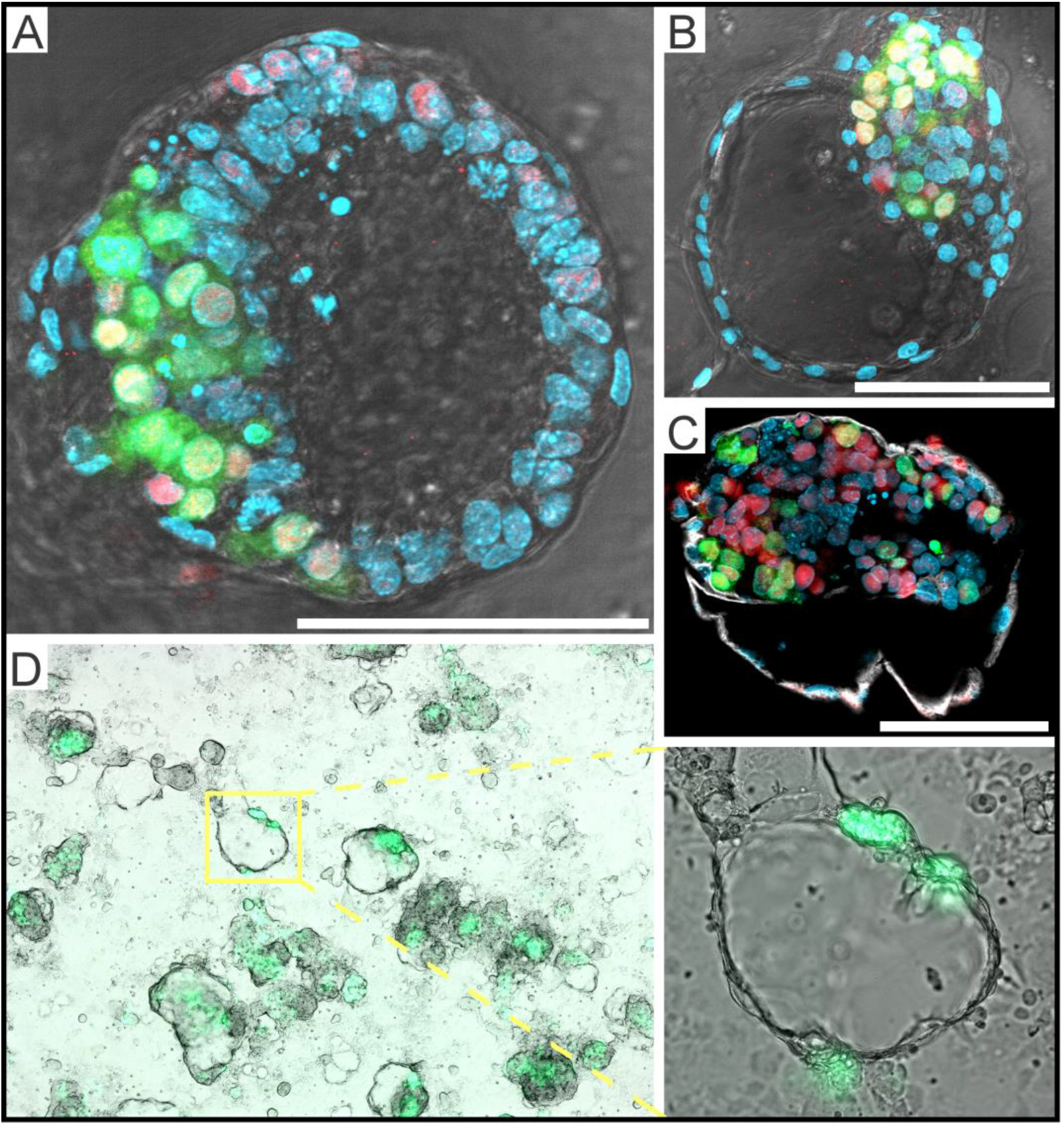
Blastocyst-Like Hemispheres Suggest Bi-Directional Potential. For All: Green = XGFP+ (Xa/Xa-GFP), Red = NANOG, Light Blue = DNA; Hoechst 33342. *All scale bars = 100 μm*. **A**) Blastocoel-like fluid filled oversized hemisphere with NANOG+XGFP+ inner cells, NANOG+XGFP-spheroid cells, and NANOG-XGFP− in flattened TE-like cells. XGFP+ cells exclusively indicate euchromatin characteristics (Figure S1A). **B**) Late BC-like hemisphere with NANOG+ cells restricted to XGFP+ cells and NANOG-XGFP− TE-like cells surrounding the fluid filled cyst. **C**) TE lineage marker positive cells (white; TROMA-I) surrounding the fluid filled hemisphere with oversized NANOG+XGFP+ polar mass. **D**) X chromosome reactivation indicated by XGFP+ cells as polar masses among fluid filled hemispheres in naive conversion experiments.

In a short time, the outer XGFP− cells completely flattened, took on morphology similar to TE, and surrounded the expanding fluid-filled cyst, wherein only the XGFP+ polar mass of the hemisphere maintained XGFP+ and NANOG+ expression (Figure 1B,C, Figure S1B, Video S1). We regularly observed tens to hundreds of such characteristic hemispheres during our previous study (Figure 1D, Figure S1A, Kime et al., 2016). Time-course reverse transcription quantitative polymerase chain reaction (RT-qPCR) experiments of naive conversion experiments revealed the induction of *Prdml(Blimpl), Prdm14, Id1, Id2, Id3, and Id4* (Figure S1C). These powerful genes broadly regulate the genome and are curiously related to the cleavage stage, early embryo, and germ line preparation (Yang et al., 2017; Hiller et al., 2010; Yamaji et al., 2008; Luna-Zurita and Bruneau, 2013; Burton et al., 2013).

The composition and organization of hemispheres drew our attention since data continued to implicate a BC: the GFP+NANOG+ polar mass of cells corresponds to the ICM where pluripotent Xa/Xa naive cells exist, and the cavity to a blastocoel. XGFP-NANOG- flattened cells had the morphology and organization of TE cells. Therefore, we checked hemispheres with an antibody specific to TROMA-I (KRT8), a well-characterized TE lineage marker. Indeed, the flattened TE morphology cells, but not XGFP+NANOG+ cells, expressed TROMA-I, and thus, the hemispheres are surrounded by XGFP-NANOG- TROMA-I+ cells (Figure 1C). Taken together, our efficient conversion of the primed state mEpiSCs to naive PSCs is concurrent with the generation of highly self-organized BC-like hemispheres.

### SMAD2/3 Signaling Inhibition and Stepwise Treatment Produces Self-Organizing Floating Blastocyst-Like 3D Structures

We tested several culture conditions to enhance the conversion efficiencies from primed to naïve PSCs. For example, adding the SMAD2/3 signaling pathway ALK5 inhibitor SB431542 had marginal effects on conversion efficiencies, yet we observed some small cell aggregates and BC-like spheres floating in the medium. We speculated that the floating spheres had BC-like properties as the hemispheres and the cell aggregates were precursors of the BC-like spheres. To yield floating BC-like structures more stably and efficiently, we optimized Phase 1 and 2 treatments with our defined supplements (Figure 2A), harvesting plate supernatants to low attachment plates on Day 6 and obtaining 5–30 floating BC-like structures by Day 7 (Figure S2A, Table S1). On Day 6 of purification, the floating structures did not stick together. Like late-hatched BCs, on Day 7 or 8, as the BC-like structures expanded, they drastically slowed growth and readily stuck together. Thus, we routinely isolated and pooled these structures at Day 7 for most downstream experiments (Figure 2B). We stained the DNA to clarify cell nuclei and found a compact ICM-like region and large flat TE-like cells surrounding a possible blastocoel similar to the hemispheres (Figure S2B). From these observations and more hereafter, we termed these structures induced blastocysts (iBCs).

**FIGURE 2:**
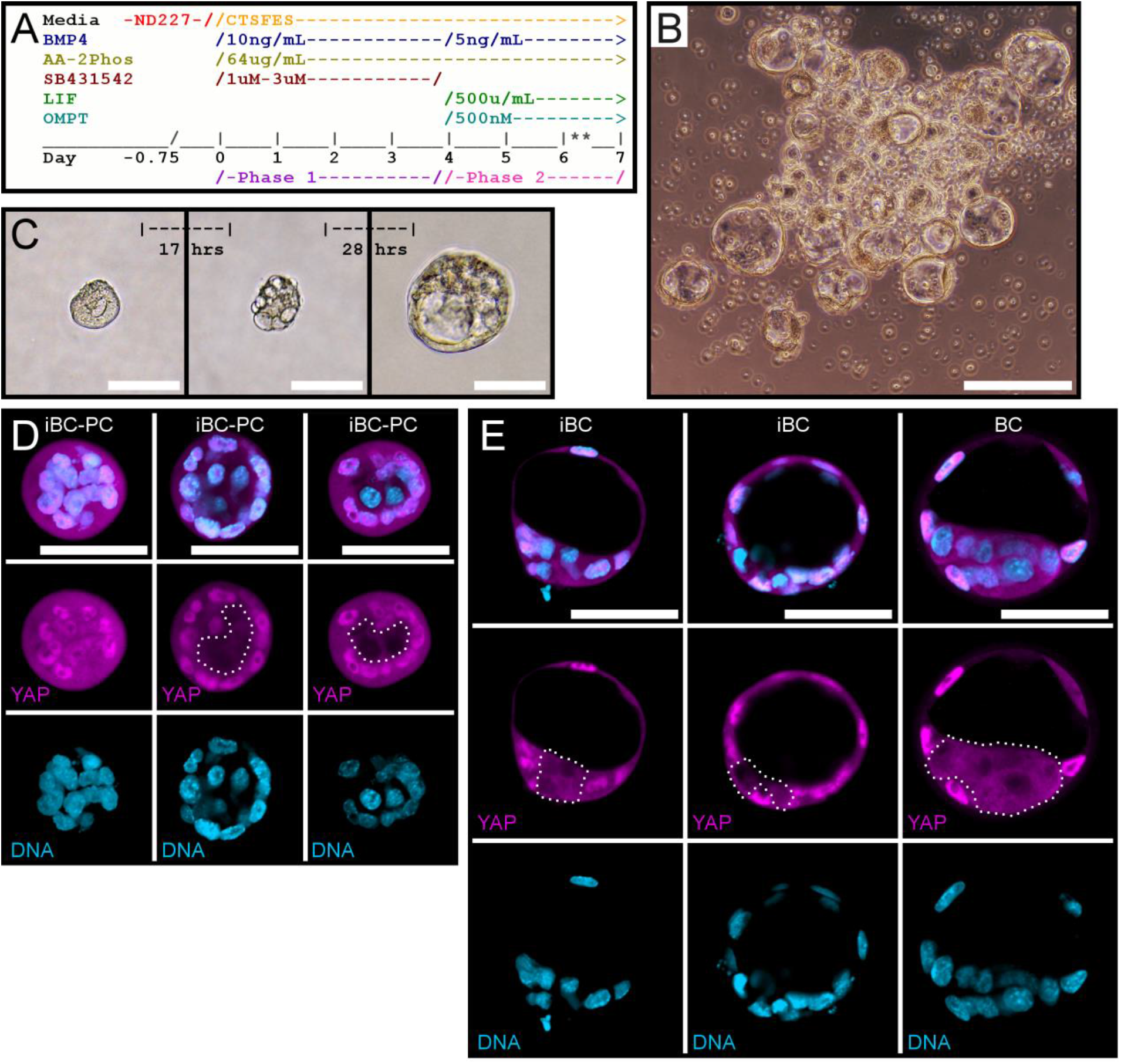
Defined Conditions Release Early Embryo-Like iBC-PCs and Polarizing iBCs to Suspension. **A**) Two-phase iBC induction media timing to induce mEpiSC to iBCs. *\*\*Supernatant iBC-PC are collected to ultra-low attachment(ULA) wells on Day 6, and high-quality BC-like iBCs are selected on Day 7 by embryo pipette*. **B**) iBCs are pooled in ULA plate for downstream experiments. *5cale bar = 200 μm*. **C**) Isolated predicted iBC-PC developing into iBC over time. *Scale bars = 100 μm*. **D**) iBC-PC stained for YAP (magenta) and DNA (light blue, Hoechst 33342). *Scale bars = 50 μm. Nuclear-excluded YAP region is outlined with dotted white line*. **E**) Early iBCs and early BCs stained for YAP (magenta) and DNA (light blue, Hoechst 33342). *Scale bars = 50 μm. Nuclear-excluded YAP region is outlined with dotted white line*.

Next, to ask if iBCs originate from a single source, we isolated the floating aggregate spheres that visibly lacked any particular polarity on Day 5.5 of induction (Figure 2C, left panel). The isolated spheres were individually cultured and regularly observed. Most grew and changed morphology: some developed into iBCs (Figure 2C). In some occasions, the isolated sphere appeared to change in the first 17 hours, resulting in mostly large round cells among a few cell types. In the next 28 hours, the cells divided and grew to further change morphology, forming an apparent cavity and became an iBC (Figure 2C). Prolonged culture thereafter resulted in paused growth and slightly reduced size of the iBC structure. These observations led us to speculate that stepwise treatment of PSC culture *in-vitro* induces floating aggregates as iBC precursors (iBC-PC) that morphologically develop and expand as iBCs.

To examine early polarity and inner/outer cell likeness, we collected iBC-PCs for immunofluorescent staining of YAP, a transcription factor involved in the first positional information and differentiation of the outer and inner cells of the early mouse embryo (Nishioka et al., 2009, Bedzhov et al., 2014). Similar to late cleavage stage non-polarized and early embryos (Hirate et al., 2015; Nishioka et al., 2009; Yu et al., 2016), YAP distribution was homogenously cytosolic and nuclear among all cells in some iBC-PCs with stochastic nuclear positioning (Figure 2D, left). However, other iBC-PCs more closely reflected 8C/16C compacting embryos undergoing outer/inner cell polarization; YAP was excluded only from the nucleus of the inner cells (Figure 2D, middle and right). We then examined emergent early iBCs and found that YAP was excluded from the nucleus of iBC inner cells and enriched in the nucleus of iBC outer cells, similar to natural early embryos (Figure 2E, Figure 3B), although iBC putative ICMs appeared smaller. These strikingly similar distributions of YAP suggest the same molecular mechanisms and signaling pathways of early embryos are installed in iBC-PCs and early iBCs.

**FIGURE 3:**
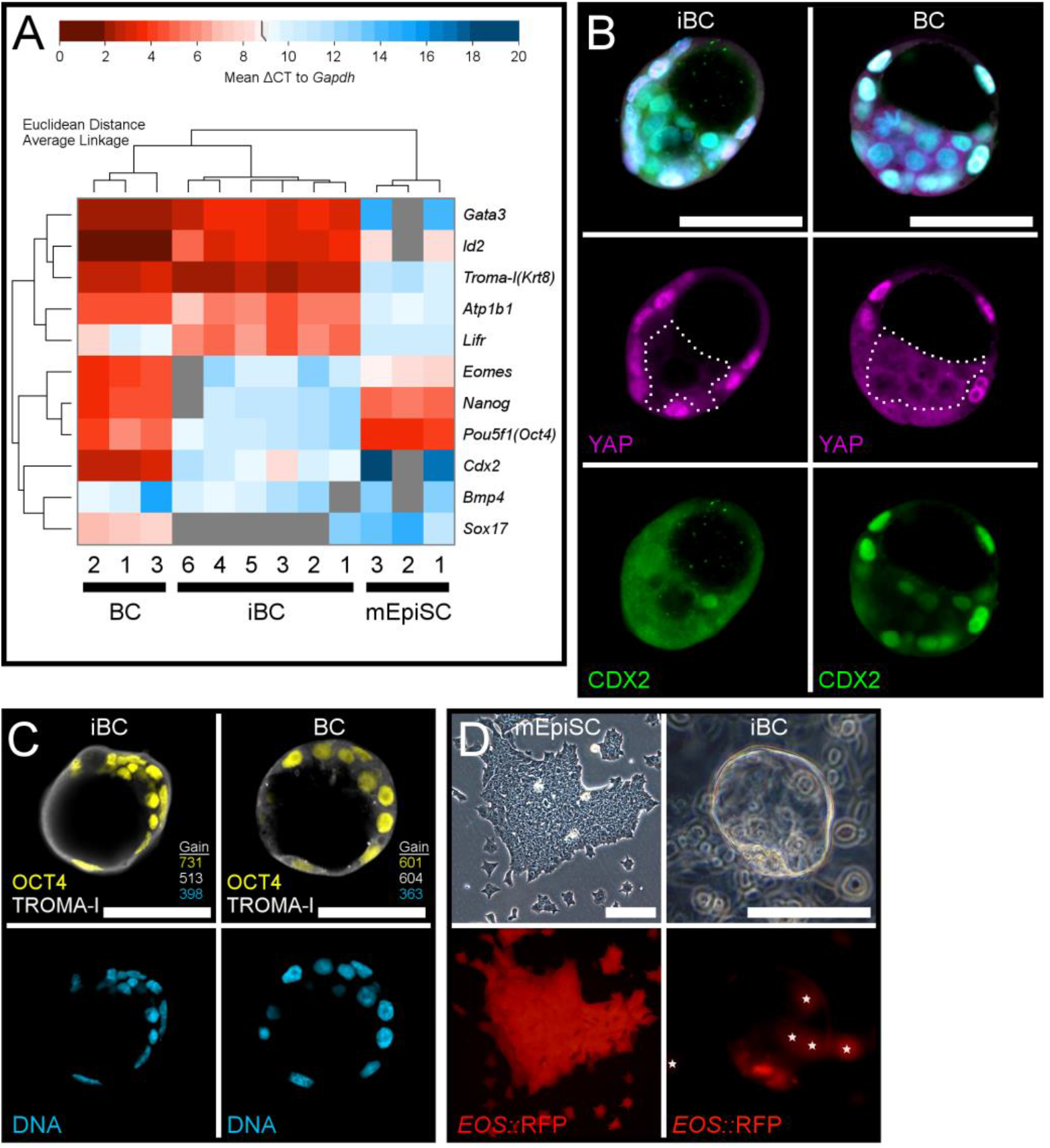
iBCs Share Many Molecular Characteristics with BCs. **A**) RT-qPCR of single BC, iBC, and mEpiSC cDNA samples, with Euclidean distance and clustering by average linkage, represented as a heat map of global ΔCT to *Gapdh*. **B**) Early iBCs and early BCs stained for YAP (magenta), CDX2 (green) and DNA (light blue, Hoechst 33342). *Scale bars = 50 μm. Nuclear-excluded YAP region is outlined with dotted white line*. **C**) Early iBC and early BC stained for for TROMA-I (white), OCT4 (yellow), and DNA (light blue, Hoechst 33342). Comparable microscopy setting detector gains have target matched colors (see methods; Table S2). *Scale bars = 50μum*. **D**) mEpiSC express live pluripotency reporter *EOS::*RFP, and late iBC above culture with EOS::RFP expression largely in the putative ICM. *White stars label out of focus *EOS::*RFP+ cells on the plate. Scale bars = 100 μm*.

BCs have an outer layer of trophoblast cells and an ICM of pluripotent cells. Thus, BCs express genes important for inducing and maintaining both lineages. To analyze BC gene expression, we extracted RNA from individually isolated BCs and iBCs, along with isolated mEpiSC colonies for a similar sized control (Figure 3A). We also sampled RNA from earlier emerging individually isolated BCs and iBCs and mEpiSC colonies accordingly (Figure S2C). RT-qPCR experiments revealed that each iBC exhibits slightly different gene expression patterns as do BCs, suggesting there is a difference in the quality, developmental timing, or both, in iBC preparations. However, overall, the key genes analyzed are expressed in iBCs at closer levels to BCs than mEpiSCs except for a few genes (Figure 3A).

Many of the genes examined that are first activated or maintained in totipotent cleavage stage cells were found strongly upregulated in iBCs to match BCs when compared with mEpiSCs (Figure 3A, Figure S2C). Interestingly, one early BC and one early iBC had comparable detectable levels of *Zscan4* (Figure S2C). In addition, the cleavage stage and naive pluripotency-related gene *Zfp42* (Rex1) was activated in early iBCs sampled (four of six), despite being undetectable in mEpiSCs (Figure S2C). Among those genes, *Atp1b1* expression was particularly striking since *Atp1b1* encodes a subunit of Na+/K+ ATPase pump, and its expression was comparable to BCs where it is essential for blastocoel formation and proper tight junctions of trophoblast cells (Hamatani et al., 2004; Madan et al., 2007). Thus, *Atp1b1* expression may be consistent with formation of a blastocoel-like fluid-filled cavity and outer layer cell tight junctions in iBCs.

Genes that are involved in the outer cell lineage induction and/or function (e.g., *Cdx2, Gata3*, and *Krt8* (Troma-I)) were also strongly induced in iBCs (Figure 3A). *Cdx2* was not equally expressed to BCs, but *Gata3* and *Krt8* were comparable. These data, with the morphology of iBC outer layer cells, suggest that a functional TE lineage may be established. In contrast, master pluripotent transcription factors *Nanog* and *Pou5f1* (Oct4) were lower in iBCs than in BCs and mEpiSCs, but still detected in many iBCs (Figure 3A, Figure S2C). *Sox2* was only detected in the *Zscan4+* iBC (Figure S2C). Furthermore, we rarely observed XGFP+ cells in iBCs (data not shown). These expressions of the pluripotency genes and XGFP are different from those in naive conversion hemispheres on the plate where XGFP+ cells expressed pluripotency genes similar to other PSCs. However, even during the naive conversion, *Nanog* is once downregulated but quickly recovers to the same level as in PSCs, dependent on LIF signaling (Kime et al 2016). Thus, early pluripotency may be induced but is not activated strongly during iBC induction.

Low-level expression of Cdx2 and Oct4 mRNA raised the question of whether iBCs correctly possess the TE lineage outer and ICM inner cell populations and are organized like BCs. To address this, we examined detection and localization of CDX2 and OCT4, along with YAP and the TE marker TROMA-I (KRT8), in early iBCs and early BCs by staining with well-characterized antibodies. The iBC inner cells downregulated CDX2 and YAP similar to BCs, but CDX2 in iBC outer cells was evenly localized and was not enriched in many nuclei (Figure 3B). These results suggest that expression and phosphorylation of CDX2 in iBCs are poorly regulated (Rings et al., 2001). Immunostaining iBCs with TROMA-I antibody revealed that outer layer TE-like flat cells are strongly positive for TROMA-I, similar to BCs (Figure 3C). Additionally, the inner cell region of the iBCs did not exhibit strong TROMA-I signals, similar to BCs, which suggested that the inner cells are not in the TE lineage. Conversely, immunostaining iBCs with OCT4 antibody clearly showed nuclear OCT4 in the inner cell region (Figure 3C; Bulut-Karslioglu et al., 2016; Ralston and Rossant, 2008). Like BCs, The nuclear signals of iBC inner cells were stronger than those in the outer layer flat cells, although iBCs have a weaker overall OCT4 signal than BCs. To investigate these differences further, we used two mainstream confocal microscopes to prepare four early iBC images as control settings to compare with early BCs. The detector gains required for a comparable image capture suggested that the strong TROMA-I signals in the outer layer cells are consistent with higher expression levels of Troma-I (Krt8) mRNA in iBCs. However, between iBCs and BCs, the detector gain difference for OCT4 signal was less than that of Pou5f1 (Oct4) mRNA, suggesting post-transcriptional regulation of Oct4 (Figure 3A,C, Figure S2C, Table S2).

At last, to examine Oct4 function as a transcription factor, we used the *EOS-S(4+)* synthetic live pluripotency reporter from the Sox2 genetic element *Srr2* to drive a red fluorescent protein (RFP, *EOS::*RFP). *Srr2* elements require a heterodimer of pluripotency transcription factors OCT4 and SOX2 to activate (Hotta et al., 2009; Tomioka et al., 2002). We introduced the reporter construct into mEpiSCs and established recombinant *EOS::*RFP mEpiSCs (Figure 3D). As expected, mEpiSCs were strongly *EOS::*RFP positive, and the RFP was not detected in differentiated cells, as reported (Figure 3D, Figure S2E; Hotta et al 2009; Tomioka et al 2002). We generated iBCs from *EOS::*RFP mEpiSCs, and the resulting iBCs often exhibited RFP signals (Figure 3D). Importantly, the RFP signals were stronger in the putative ICM than outer cells, suggesting OCT4 and SOX2 in the inner cell regions of iBCs better activate or maintain *EOS::*RFP expression. Yields from three iBC production wells were 14, 20, and 27 iBCs, and among those iBCs, 12, 13, and 20 had detectable *EOS::*RFP+ (Figure S2F). These data suggest iBCs have an ICM-like region where some pluripotency transcription network is operative.

Thus, iBC-PCs and iBCs exhibit several unique features that are common with preimplantation embryos at morphological and molecular levels. We concluded that the stepwise manipulation of signaling pathways triggered, to some extent, a dynamic self-organizing event reminiscent of the first polarization, differentiation, and morphogenesis of early embryonic development.

### Reproducibility of iBC Generation

Our various XGFP mEpiSC reporter sub-lines performed similarly throughout iBC induction. Two published mEpiSC lines reacted similarly throughout and produced iBCs with lower yields (Figure S2G; Tesar et al., 2007; Ohtsuka et al., 2012). Another mEpiSC line with apparent cell culture characteristic differences failed completely (Parchem et al., 2014). Therefore, iBC generation should be possible with many but not all mEpiSC lines.

### Purified iBCs Implant, Induce Decidualization, and Grow in Pseudopregnant Mice

#### Implantation/Decidualization Rates of iBCs

The characteristic similarities between iBCs and BCs led us to examine implantation-competence and developmental potency of iBCs *in-utero*. We collected iBC-PCs at Day 6 or 7 of induction and allowed them about 18 hours to expand toward iBCs that were purified with an embryo pipette by visual assessment and loosely pooled. Purified iBCs were transferred to the uterus horn of sterile-male bred pseudopregnant mice, and positive control BCs were transferred separately; we also transferred large mEpiSC colony clusters and embryoid bodies (EBs) prepared from mEpiSCs as controls. We dissected the transplanted mice between E5.5 and E7.5 and found that transferring iBCs induced deciduae in the pseudopregnant mice while mEpiSC clusters and EBs failed (Figure 4A,B). Deciduae induced by iBC transfer were similar in focal morphology to BC-induced deciduae although they were, on average, smaller in size than those from BC transfers. Importantly, many iBC-induced deciduae recruited large maternal blood vessels seen in the uterus, and sectioning showed red brown color in the decidua basalis region, similar to those from BCs, indicating significant blood supply from the mother (Figure 4B). These observations clearly show that iBCs induce decidualization of the uterus and recruit the maternal blood supply, which result from iBC implantation. Still, the smaller sized deciduae from iBCs suggested postimplantation developmental delay or resorption.

**FIGURE 4:**
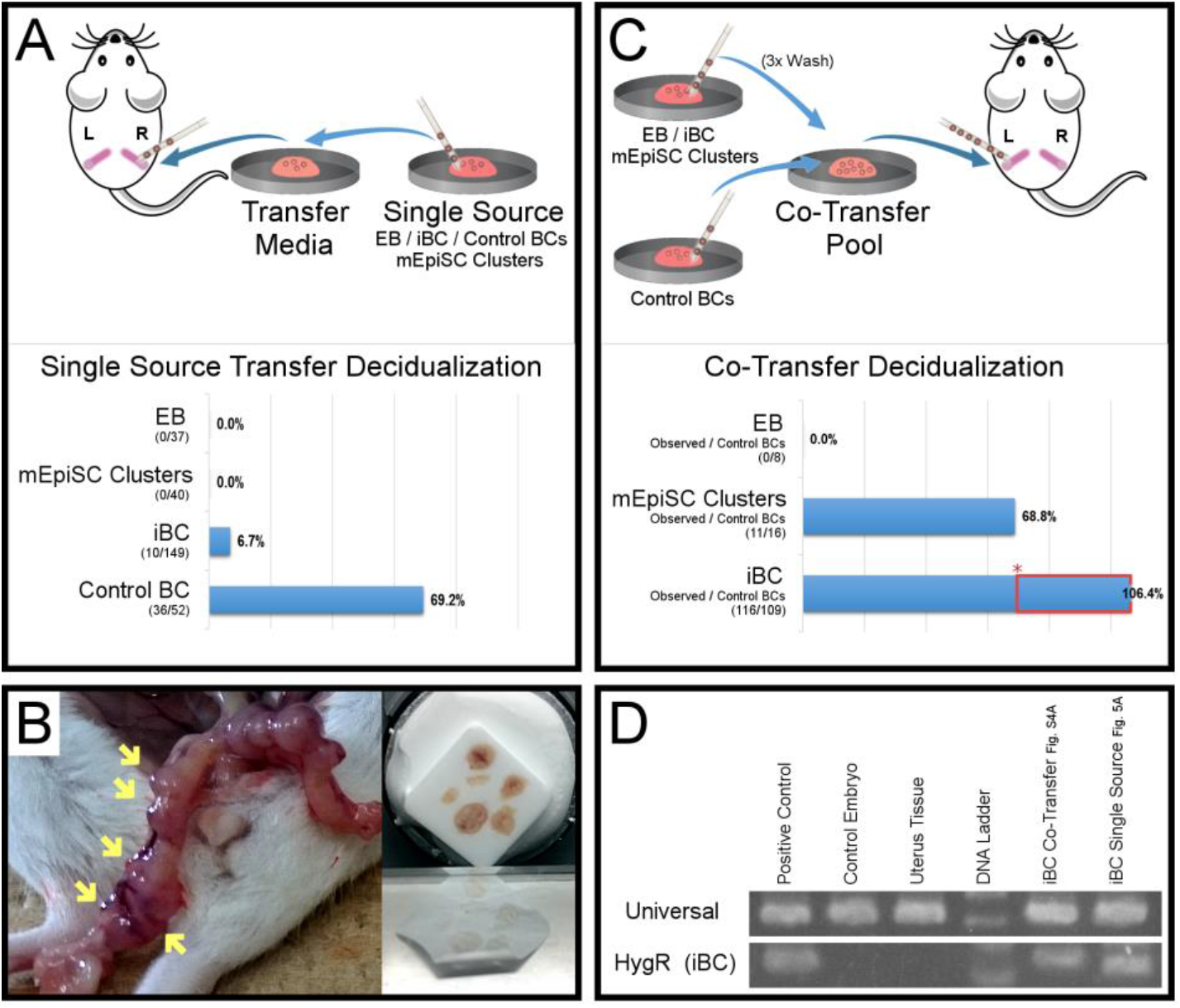
iBC Uterus Transfer Decidualization in Pseudopregnant Mice. *Embryo pipettes indicate when a new pipette was used*. **A**) Single source uterus transfer experiment diagram. Observed deciduae in uterus horns with respect to EB, mEpiSC clusters, iBC, or control BC single source uterus transfers. **B**) Uterus horn of mouse with iBC implanted deciduae (left, yellow arrows), prepared for cryosection (right). **C**) EB or mEpiSC clusters or iBC co-transfer with BCs uterus transfer experiment diagram. Observed deciduae from EB co-transfer, mEpiSC Cluster co-transfer, and iBC co-transfer. * = control BC decidualization rate (69.2%, Figure 4A); Red box indicates decidualization gained from iBCs.) **D**) LCM genomic DNA PCR test for mouse genomic DNA universal and iBC-specific hygromycin resistance. LCM samples from figure H&E slides are shown in Figure S4A.

By counting transferred iBCs or BCs and the resulting deciduae obtained, we calculated frequencies of decidualization (Table S3). iBCs and BCs induced deciduae at 6.7% (10/149) and 69.2% (36/52), respectively (Figure 4A). Thus, iBCs alone induce decidualization but less frequently than BCs. The lower frequency by iBCs may be due to different iBC developmental size and timing among a heterogeneity in quality (Figure 2, Figure 3, Figure S2).

Co-transferring control embryos that easily implant improves implantation rates of difficult embryos in assisted reproductive settings (Mochida et al., 2014). We therefore transferred iBCs in co-transfer with control BCs (Figure 4C). In total, we transferred 186 iBCs along with 109 BCs to numerous uteri, and upon dissection, we frequently observed more deciduae than the number of control BCs, implicating iBCs as the source of increased implantation. Among *all* co-transfer experiments, the total number of 116 observed deciduae divided by the total number of 109 control BCs would mean an impossible implantation rate of 106.4% (116/109) if control BCs were the only source (Figure 4C, Table S3). We note that, since control BCs transferred alone implanted at 69.2% (Figure 4A) and others reported control BC implantation rates of 43-44% (Bulut-Karslioglu et al., 2016), iBCs should be the source of the 37.2% difference, or increase the implantation of control BCs, or a combination therein (Figure 4C). Yet even if we assumed 100% success from all BCs, we performed 26 total iBC+BC co-transfer experiments, and 15 of those experiments (57.7%) induced more focal deciduae than the number of BCs (Table S3). Given that co-transfer is more robust, we also examined co-transfer of mEpiSC clusters or EBs from mEpiSCs along with BCs. We found no deciduae from the EB+BC co-transfers, meaning the size or cells of EBs might impair the implantation process (Figure 4C). mEpiSC clusters with BCs proved to be a better control co-transfer experiment, allowing for the likely 68.8% implantation of control BCs similar to control BCs alone (Figure 4A,C).

iBC implantation and decidualization in co-transfer with BC are higher than as a single source. To confirm the origin of the deciduae from both types of experiments, we obtained genomic DNA from some deciduae cryosections by laser capture microdissection (LCM) and amplified a transgenic DNA region specific to cells that generate iBCs (Figure 4D, Figure S2D, Figure S4A). This analysis confirmed that iBCs from both the single source and co-transfer experiments induced deciduae with detectable tissue at the proper location for natural embryos (Figure 4B). We therefore recognize that co-transferring BCs with iBCs enhances the ability of iBCs to induce decidualization and may prove more useful in later studies.

#### Cryosection and Analysis of Implanted iBC-Derived Tissues

We performed hematoxylin and eosin (H&E) staining and immunohistochemistry (IHC) on cryosections of dissected deciduae from E7.5 iBC single source transfer experiments and compared them to control embryos at E7.5 and E6.5 (Figure 5, Figure S4B). Similar to control deciduae, iBC-induced deciduae were surrounded by uterine tissue and had distinct sub-regions; among those, the decidua basalis showed vascular sinus foldings and red blood cells, indicating maternal blood supply (Figure 5, Figure S3, Figure S4A,B). Sections often showed distinct disfigured tissues in the presumptive embryonic region with surrounding ExEm-like cells and internal small dark stained cells resembling the embryonic portion (Figure 5A,C, Figure S3, Figure S4A). We tested proximal cryosections of the same deciduae for ExEm lineage TROMA-I and found that the surrounding iBC-derived ExEm-like tissues were TROMA-I+ and had invaded the deciduae, growing to a total size similar to or in excess of a control E6.5 embryo (Figure 5B). The cryosections showed tissues resembling a retracting parietal yolk sac cavity, a degrading putative Reichert’s membrane, and TROMA-I+ cells surrounding internal TROMA-I- cells that we speculated to be Em portion cells based on location and H&E stain characteristics (Figure 5, Figure S3, Figure S4A). While iBC-derived E7.5 tissues were larger than E6.5 control embryo tissues, many cells appeared pycnotic and lacked a healthy appearance, and were collectively smaller than a E7.5 control embryo (Figure 4B; Gardner and Johnson, 1972).

**FIGURE 5:**
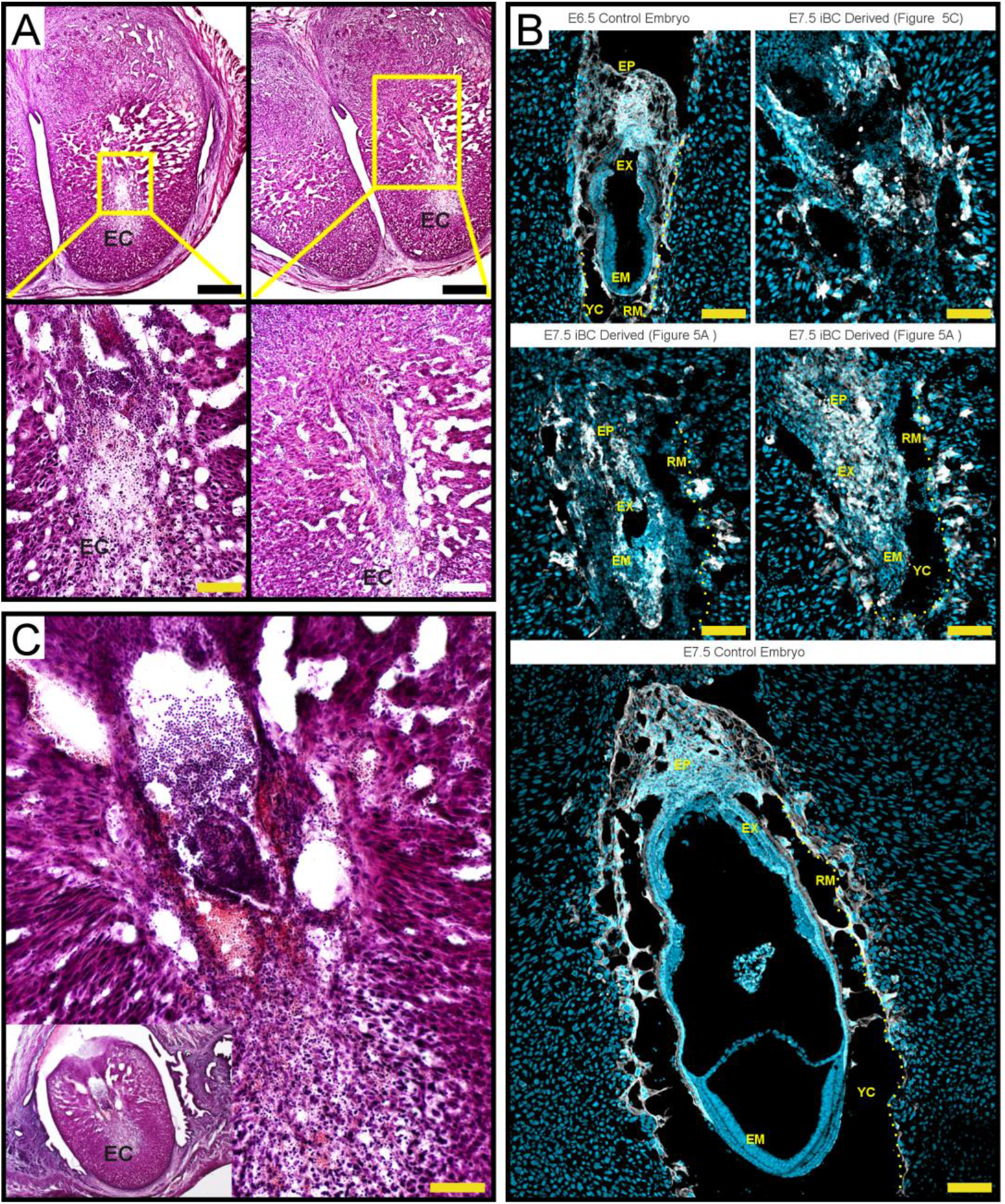
iBCs Implant and Partially Develop Before Resorption. *EC, Embryonic Cavity* **A)** H&E-stained proximal cryosections of deciduae from iBC single source transfer. Higher magnification is indicated and shows iBC-derived tissue resembling large cell masses of resorbing tissues. *Scale Bars: Upper panels = 500 μm, Lower Left = 100 μm, and Lower Right = 200 μm*. **B)** Cryosection IHC for ExEm TROMA-I (white), and DNA (light blue; Hoechst 33342). E6.5 and E7.5 control embryos show healthy size and structure. E7.5 iBC-derived tissues from cryosections proximal to Figure 5A and Figure 5C are labeled. *EP, ectoplacental cone; EX, extraembryonic portion; EM, embryonic portion; YC, yolk sac cavity; RM & dotted line, Reichert’s Membrane labeled on one side for clarity. Scale bars = 100 μm*. **C)** H&E stained decidua from iBC single source transfer section shows high presence of immune cells resorbing a mass of cells with ExEm-like and Em-like stain and morphology. *Scale bar = 100 μm*.

To further examine the development of iBC-derived ExEm tissues in implanted deciduae, we performed IHC for ExEm markers trophoblast-specific protein alpha (TPBPA) and placental lactogen 1 (PL-I). TPBPA is expressed in ectoplacental cone, spongiotrophoblasts and precursors of trophoblast giant cells (TGCs) that originate from TE (Simmons and Cross, 2005; Simmons et al., 2007). TPBPA was detected in the larger cells surrounding the embryonic cavity in both iBC and BC-implanted deciduae at E6.5 (Figure S4C). At the same time, PL-I expressing cells characteristic of parietal TGCs lined the embryonic cavity in iBC deciduae similar to the control embryo where it was also detected on the visceral endoderm (Figure S4C; Chen et al., 2016; Peng et al., 2015; Simmons and Cross, 2005; Simmons et al., 2007; Screen et al., 2008). Some TPBPA+ or PL-I+ cells had larger more brightly stained nuclei and scattered far from the cavity, suggesting they are polyploid scattering TGCs (Figure S4C).

#### Implanted iBCs May Develop Briefly Before Resorption

We noticed that iBC-implanted deciduae were often variably smaller than control deciduae of the same timing. Histology frequently showed evidence of a retracting post-implantation embryonic cavity (Figure 5A,C, Figure S4). We observed blood cells within the blood sinuses and around the iBC-derived implanted tissues (Figure S3), but the high presence of lymphoid and myeloid cells around the disfigured pycnotic tissues indicated the embryo resorption process (Figure 5C, Figure S3; Cossée et al., 2000; Flores et al., 2014). Still, upon closer observation, iBC-derived non-decidual tissues were markedly diverse and had morphology and localization similar to invading trophoblasts, ectoplacental cone, ExEm portion, Em portion, and yolk sac cavity, when compared to a previous report of natural resorbing embryos (Figure 4, Figure S3; Cossée et al., 2000).

These data show that some iBCs are functionally competent to implant, induce decidualization, and grow to greater cell numbers while developing over several days. Implanted iBC tissues displayed varied natural analogous characteristics while providing distinct evidence of ExEm tissue differentiation surrounding internal cells resembling the Em lineage, and in total, apparently following a natural progression of embryonic resorption.

### Further Characterization of the iBC Generation Process

#### Establishing Pluripotency Is Insufficient in iBCs, Yet Possible in Outgrowths

Mouse embryonic stem (ES) cells are a naive PSC-derived from the ICM of BCs. In addition, trophoblast stem (TS) cells can be established from BCs using different conditions. To determine if ES-like or TS-like cells could be derived from iBCs, we isolated iBCs/iBC-PCs with XGFP and EO5/:D2nRFP dual reporters and plated them on feeder cells.

In ES cell derivation conditions (Czechanski et al., 2014), outgrowths proliferated, and some cells expressed XGFP and *EOS::*D2nRFP (Figure S5A). These cells could be passaged and enriched like naive ES cells and were comparable to naive ES cells when stained for OCT4, NANOG, and YAP (Figure 6A). We also examined expression of several important pluripotency and early embryonic genes with RT-qPCR and found that iBC/iBC-PC derived cells were generally comparable to ES cells (Figure 6B). Interestingly, iBC/iBC-PC-derived cells expressed notably higher levels of *Zscan4* and *Zfp42* (Rex1).

**FIGURE 6:**
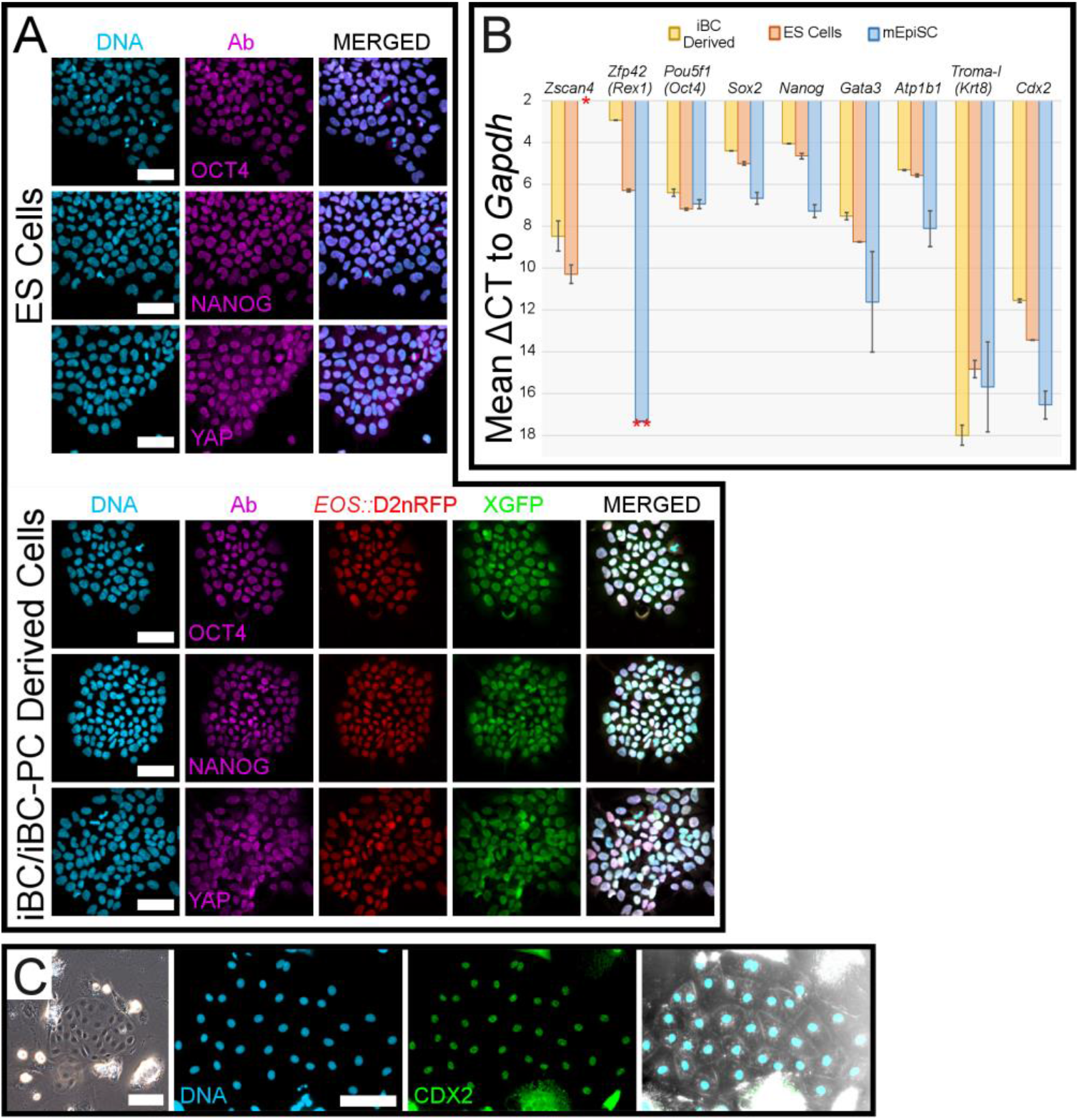
iBC/iBC-PC-Derived Outgrowths Suggest *In-Vitro* Bi-Directional Potential. **A**) Mouse ES cells and iBC/iBC-PC-derived ES-like cells stained for OCT4, NANOG, or YAP (magenta), and DNA (light blue, Hoechst 33342). iBC/iBC-PC-derived ES-like cells demonstrate X chromosome reactivation (XGFP+) and express *EO5::*D2nRFP. *5cale bars = 50 μm*. **B**) RT-qPCR of mouse ES cells, iBC/iBC-PC-derived ES-like cells, and mEpiSC cDNA samples, shown as mean ΔCT to *Gapdh*. Data represents two biological samples per type and all probes tested in technical triplicate. Error bars represent standard deviation between the two biological samples. * = no detectable *Zscan4* signal in mEpiSC samples. ** = one mEpiSC biological sample did not have detectable *Zfp42* (Rex1), and therefore no standard deviation. **C**) Left: Live imaging of iBC derived TE-like cells. Right: TE-like cells were stained for CDX2 (Green) and DNA (light blue, Hoechst 33342). *Channels shown separately and merged. 5cale bars = 100um*.

We asked if iBCs/iBC-PCs give rise to TE lineage progeny on feeder cells and first saw colonies of slow growing TE-like cells expressing CDX2 (Figure 6C). We also tried a defined TS cell medium to derive TS cells or their progeny (Latos and Hemberger, 2016; Ohinata and Tsukiyama, 2014), but we could not obtain stable TS cells. However, we could expand TE-like cells for a few passages in the defined condition (Figure S5B), and those cells passed through a binucleate phase that is characteristic of TE-derivative cell cultures that produce trophoblast giant cells (Figure S5B, mid panel; Ilgren, 1981). The culture of cells slowed almost to a halt of large single and binucleate cells that stained strongly but variably for the TE-lineage markers PL-I and TPBPA (Figure S5B; Awonuga et al., 2011).

These outgrowth results suggest that iBCs/iBC-PCs hold increased potential whereby additional culture time might fully establish naive pluripotency. In addition, our TS cell culture outgrowth did not establish stabilized TS cells, but yielded cells with important post-implantation ExEm markers and characteristics *in-vitro* to strengthen our cryosection IHC detection of iBC-derived implanted tissues (Figure S4C).

#### iBC Induction Requires Prdm14 and May Concomitantly Activate a Totipotency-Related Cell Program

Because iBCs coordinately differentiated evidence of a bi-directional 3D cyst expressing cleavage stage-initiated genes, such as *Atplbl*, and possibly had interim loss of pluripotency, we hypothesized that iBCs emerge from totipotent-like cells. The totipotent genome is prepared in the germ line and activated by ZGA as the zygote enters 2C and cleavage. Phase 1 of iBC induction has defined molecules involved in germ cell differentiation (Chen et al., 2012; Hikabe et al., 2016; Yamaji et al., 2008; Yang et al., 2017), and Phase 2 has defined molecules that induce naive pluripotency and TE lineage transdifferentiation (Figure 2A; Bao et al., 2009; Hayashi et al., 2010; Kime et al., 2016). In this sense, we looked to Prdm14, a major gene regulatory factor shared in the germ line and early embryo (Nakaki and Saitou, 2014; Hackett et al., 2017). RT-qPCR of mEpiSCs showed very low detection of *Prdm14* in large samples, and *Prdm14* was undetectable in small isolated colonies; however, in experiments for isolated BC and iBC samples, some iBCs expressed *Prdm14* at significant or comparable levels to BCs (Figure 7A), suggesting that this important transcription factor is induced in the iBC process.

**FIGURE 7:**
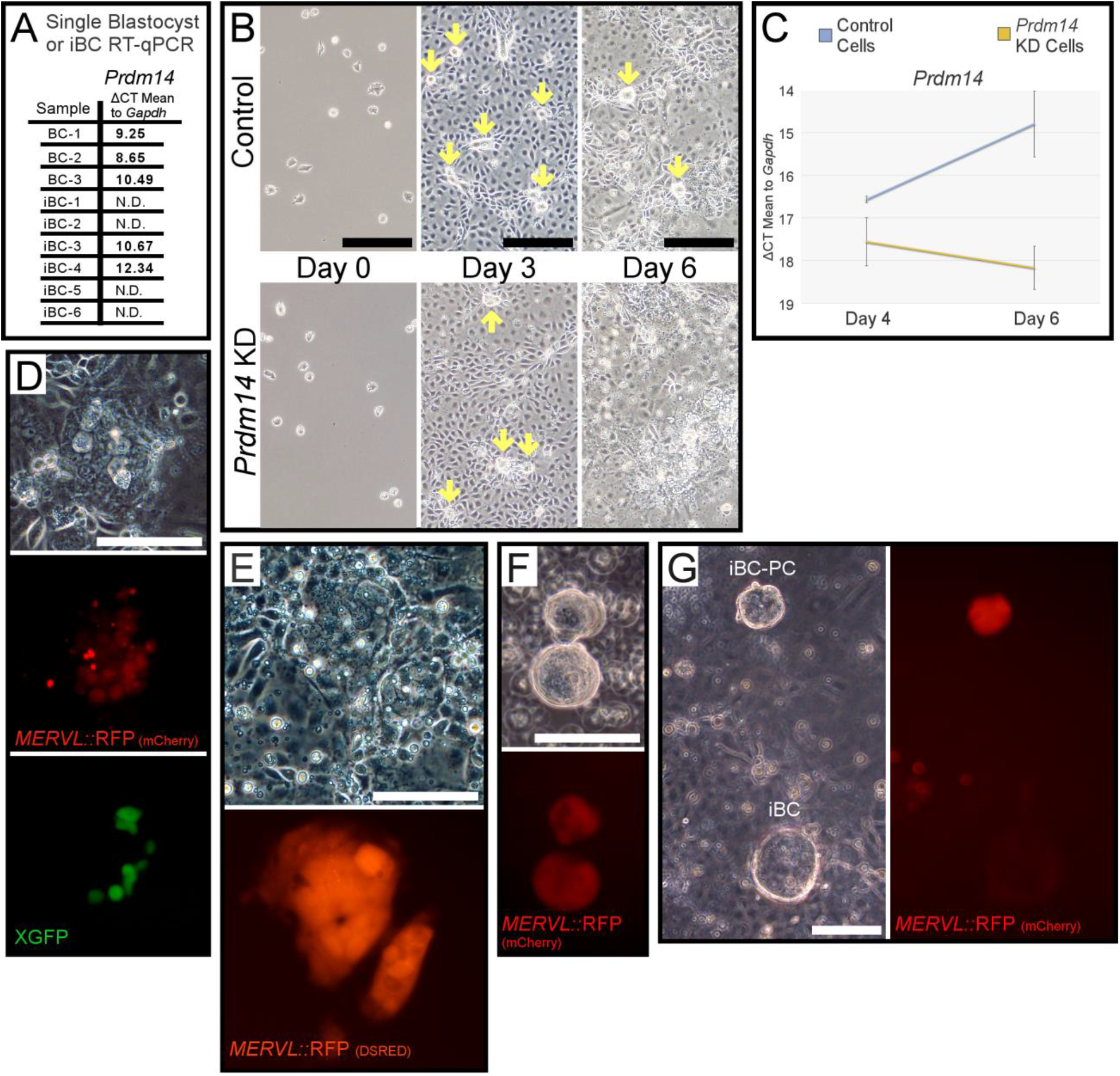
iBC Generation Requires Prdm14 and Activates MERVL Reporter in iBC-PCs. **A**) Single isolated BC and iBC RT-qPCR for *Prdm14*. **B**) Control and *Prdm14* KD mEpiSC are plated for iBC induction. Loci that originate iBC-PC are initiated in both experiments by Day 3 (yellow arrows). Control cells maintain iBC-PC induction through Day 6 (yellow arrows) and *Prdm14* KD cells abort iBC-PCs among cell debris. *Scale bars = 200um*. **C**) RT-qPCR of control and *Prdm14* KD cell plate cDNA samples for *Prdm14* in iBC generation, shown as mean ΔCT to *Gapdh*. *Error bars represent standard deviation from technical triplicate*. **D**) iBC induction Day 6 colocalized expression of *MERVL::*RFP and XGFP+ reporters. *Scale bar = 100 μm*. **E**) After iBC are collected, iBC generation plate on Day 7 retained some larger *MERVL::*RFP+ cells with cleavage stage cell-like morphology. *Scale bar = 100um*. **F**) Live fluorescent image of iBC-PCs expressing *MERVL::*RFP in ULA plate on Day 6. *Scale bar = 100 μm*. **G**) Live fluorescent image of *MERVL::*RFP expressed strongly in iBC-PC yet poorly detected in emergent iBC, seen in culture on Day 8. *Scale bar = 100 μm*.

We tested our constitutive shRNA *Prdm14* knockdown(KD) mEpiSCs and found with daily microscopy that the experiment began similar to control cells and initiated the compacted iBC-PC/iBC originating loci (Figure 7B). However, by Day 6, the iBC-PCs that may become iBCs were nearly completely aborted, and the peripheral cells appear to degrade (Figure 7B). Supernatants collected from all iBC experiments with *Prdm14* KD cells failed to yield favorable iBCs and had increased cell debris. For reference, when control cell iBC-PC were not harvested by agitation to supernatant on Day 6, they still differentiated as gently attached expanded iBC above such concentrated cell loci (Figure 3D, Figure 7G).

To examine *Prdm14* expression in control and *Prdm14* KD cells, we collected mRNA from cell populations at the end of Phase 1 on Day 4, when the cultures appear to perform similarly, and on Day 6, immediately before iBC-PCs are usually harvested or lost in the *Prdm14* KD cell population (Figure 7B). Compared to the detection of *Gapdh*, control cell iBC induction populations had notable *Prdm14* expression by Day 4, and further increased by Day 6. *Prdm14* KD cells showed significantly reduced *Prdm14* expression on Day 4 and a much lower proportional expression by Day 6 (Figure 7C). The lower detection of *Prdm14* in Prdm14 KD cells on Day 6 suggests that the lost iBC-PCs required *Prdm14* and that iBC-PC might represent most of the detectable *Prdm14* at that time (Figure 7B,C).

To further elucidate a possible relationship between totipotency and iBC generation, we cloned the well-studied 2C *MERVL* live totipotency-related reporter to drive an RFP(*MERVL::*RFP) in XGFP mEpiSCs to operate dual reporters (Figure S2D). These reporters were undetectable in mEpiSCs in agreement with previous reports (Bao et al., 2009; Macfarlan et al., 2012; Wu et al., 2017). On Days 5-6 of iBC induction, we observed some of the characteristic loci where iBC-PC originate had *MERVL::*RFP+ cells, and many cells expressed XGFP, suggesting X chromosome reactivation to Xa/Xa (Figure 7D). We speculated that cells with dual reporter activation implicates ZGA mechanisms since both are reported characteristics of 2C cleavage stage cells (Figure 7D; Monk and Harper, 1979; Okamoto et al., 2004, Wu et al., 2017). Interestingly, when iBC-PC were harvested, some remaining attached *MERVL::*RFP+ cells variably lost XGFP expression and often became larger and more rounded with cleavage stage-like cell morphology (Figure 7E). Furthermore, despite the low overall frequency of *MERVL*+ reporter cell loci on the plate at Day 6, many of the iBC-PCs harvested by agitation were composed of *MERVL::*RFP+ cells (Figure 7F). *MERVL::*RFP expression in iBC-PC was usually weaker than the *MERVL::*RFP+ cells seen on the plate (Figure 7D,E,F,G). Also, XGFP was generally not observed in *MERVL::*RFP+ collected iBC-PCs, suggesting that both reporters were down regulated at that critical stage similar to compacting 8C/16C embryos that precede BCs. Strengthening this observation, emergent iBCs had far reduced detectable *MERVL::*RFP (Figure 7G). Of further interest, *MERVL::*RFP+ and *MERVL::*RFP+/XGFP+ cells were variably maintained on the plate for several days in Phase 2 media after iBC-PC harvest.

Collectively, our results indicated that the iBC production process required *Prdm14*. *MERVL::*RFP+/XGFP+ subpopulations preceded and reported from the characteristic loci where *Prdm14* dependent iBC-PC emerged, and many harvested iBC-PCs expressed the *MERVL::*RFP reporter. These data show an unknown intermediate role for Prdm14-dependent iBC formation among plate loci where totipotent-cell characteristics may be differentially but strongly induced. Since *Prdm14* KD compromised the entire iBC formation via abortive loss of iBC-PCs, and both the TE-like and putative ICM-like cells were lost, Prdm14 may be key to improved induction of totipotent 2C-like cells or the specification of lineages in iBCs.

## DISCUSSION

This study showed that sequential treatment of mouse PSC culture with defined molecules reproducibly induces BC-like 3D structures with several distinct features of BCs. Remarkably, the structures emerged from floating small cell clumps resembling denuded 8C/16C-compact embryos wherein cells lack morphological differences or polarity. As such, differentiation to the iBC structure is emergent. iBCs exhibited signs of implantation when transplanted into pseudopregnant surrogates, indicating implantation-competence. In previous work of mouse PSC-derived oogenesis, rare BC-like structures from 40+ day long-term differentiation experiments were partially described (Hübner et al., 2003). However, whether those structures were developmentally competent *in-utero* is unknown. Furthermore, using TS cells, PSCs and PSC-derived bi-directionally contributing cells has never been demonstrated to contribute to animal development in transplanted pseudopregnant mothers without donor cells or chimerism for support (Macfarlan et al., 2012; Yang et al., 2017). Therefore, generation of fully functional iBCs, which give rise to newborn animals in an isogenic setting, may uncover a maximum differentiation potential of PSCs.

To our knowledge, this is the first demonstration that PSC culture can generate the 3D architecture with cellular materials and implantation-competence resembling BCs. Until now, only implantation-competent BCs or their trophoblasts (Gardner and Johnson, 1972), chimeras thereof, or specific melanoma cells were reported to induce deciduae in sterile-male bred pseudopregnant mice (Wilson, 1963). A related field of uterine environment study involves deliberate uterus disruption often combined with injected progesterone and estrogen hormone treatments to induce deciduomas (Herington and Bany, 2007; Lee et al., 2007). Deciduomas are composed of homogenous decidual cells, and focal deciduomas that better resemble individual natural deciduae requires concanavalin A-coated Sepharose beads (Herington et al., 2009). Unlike deciduomas, our iBC-induced deciduae rely exclusively on sterile-male bred pseudopregnant surrogates and have correct positional implanted tissues in focal deciduae that resemble natural deciduae or deciduae from BC-derived trophoblast vesicles (Gardner and Johnson, 1972). To exclude the possibility of deciduomas, we molecularly characterized iBC-derived implanted tissues and emphasize that we do not use the materials or methods required to produce deciduoma. In our experiments, co-transfer with control embryos may greatly improve the implantation of the iBC, as with difficult mouse strains (Mochida et al., 2014), yet iBC alone demonstrate implantation-competence.

### Molecular Considerations in the iBC Process

We showed that YAP localization implicates non-polarized and polarized iBC-PC, and polarized early iBCs, that are critically similar to early embryos. Intermediate MERVL reporter and comparable *Zscan4* activation furthers that prospect. Signaling inputs that we provided during iBC production may mimic developmental cues of embryogenesis. Using a synthetic LPA (OMPT) in our cocktail may be striking because LPA treated BCs exhibit enhanced embryogenesis by activating YAP *in-vitro* and *in-utero* (Yu et al., 2016).

iBCs expressed *Cdx2* at a lower level than BCs and had some nuclear CDX2 localization while most iBC outer cells retained cytosolic CDX2. During early development, Cdx2 is preferentially upregulated around the 8C stage to specify committed outer cells (Strumpf et al., 2005; Ralston and Rossant, 2008). Cdx2 is crucial for development since Cdx2-deficient BCs cannot implant in the uterus despite having functional pluripotent cells in the ICM (Meissner and Jaenisch, 2006). We found iBCs exhibit a TROMA-I+ and nuclear-enriched YAP outer layer and a blastocoel-like cavity and are implantation-competent. Additionally, TROMA-I+ cells from transplanted iBCs grew well, invaded the uterus to decidua reaction, and developed different morphologies and detectable markers (PL-I, TPBPA), depending on their positions in the embryonic cavity. These results further indicate the proliferation and differentiation capacity of the iBC outer cells within the TE lineage after implantation despite weak iBC Cdx2 characteristics. Additionally, post-implantation proliferation of trophoblast progeny depends on ICM-derived tissues in normal development, which suggests that some of the larger more developed iBC-derived implanted tissue may have been helped by cells from the iBC ICM-like region (Gardner and Johnson, 1972, 1975; Rossant and Ofer, 1977; Simmons and Cross, 2005).

iBCs plated on feeder cells produced TE-like colonies uniformly expressing CDX2 protein, and when grown in defined TE cell culture conditions, the iBC/iBC-PC-derived cells were characteristically similar to dissociated cultured TE derivatives. Thus, Cdx2 expression in iBCs may be sufficient to induce functional TE lineage cells that enable iBCs to implant and develop for several days. However, we observed that iBCs had uneven implantation, implicating molecular pathways that interface between the TE-like cells and receptive uterus are incorrect. Optimizing our regimen based on known conditions that induce trophoblasts from PSCs may improve Cdx2 expression to correct abnormalities (Hayashi et al., 2010).

Our results show that iBCs have a putative ICM with OCT4 and nuclear-excluded YAP and downregulated both CDX2 and TROMA-I protein. Notably, the exclusion of nuclear YAP is a characteristic of pluripotent cells in the ICM of BCs that is critically different from *in-vitro* cultured mouse pluripotent ES cells that have nuclear-enriched YAP (Tamm et al., 2011; Figure 6A). We speculate that the putative ICM in iBCs became the central TROMA-I- cells we observed in iBC-derived cryosections. However, the key pluripotent transcription factor Oct4 mRNA was expressed at lower levels in iBCs than in BCs. Since precise expression of Oct4 is crucial for establishing and/or maintaining pluripotency, lower expression suggests suboptimal re-establishment of pluripotency in iBCs (Niwa et al., 2000). This may explain why iBC-derived post-implantation proliferation and development eventually delayed or ceased (Gardner and Johnson, 1972). Additionally, we rarely observed Xi-GFP reactivation in iBCs, consistent with the lower expression of *Nanog* (Silva et al., 2009). However, we could establish ES cell-like cells from iBCs/iBC-PCs in naive PSC derivation conditions, suggesting that authentic pluripotency could be reestablished in altered conditions. We therefore speculate that minor critical adjustments to iBC generation conditions may improve intermediate iBC-PC and iBC cell states.

The findings of the insufficient pluripotency in iBCs are in stark contrast to what we observed during the hemisphere formation experiments where naive pluripotency was robustly established (Kime et al., 2016). In that study, we showed that BMP4 signaling, with LIF and ascorbic acid, greatly increased *Prdm14* expression during conversion of mEpiSCs to the naive state (Kime et al., 2016); but we also measured the induction of Prdm1(Blimp1) and *Id* gene family mRNAs, which we reported here. SMAD2/3 signaling inhibition stimulates BMP induction, and BMP4 can replace serum in naive PSC culture by inducing Id genes toward self-renewal (Ying et al., 2003). BMP signaling and these critical Prdm and Id family genes are shared among germ cell development and cleavage through preimplantation embryonic development (Yang et al., 2017; Hiller et al., 2010; Yamaji et al., 2008). Interestingly, *Id2* is comparably expressed in outer and inner lineage early embryonic cells (Ying et al., 2003; Tang et al., 2010; Wu et al., 2016). One considerable difference between our hemisphere/naive conversion and iBC generation is in the induction regimes. SMAD2/3 signaling inhibition may be necessary to generate iBCs, but SMAD2 specific inhibition induces TE and germ cell differentiation while suppressing pluripotency expression via Nanog inhibition (Chen et al., 2012; Sakaki-Yumoto et al., 2013). We anticipate that SMAD2/3 signaling inhibition may require further adjustment to achieve sufficient pluripotency in iBCs and note that a necessary adjustment of SB431542 concentration was mEpiSC line specific.

*Prdm14* KD experiments suggest that Prdm14 has a pivotal role during iBC induction although the exact mechanisms are unknown. In the iBC system, *Prdm14* was greatly enriched by Day 4, prior to LIF in Phase 2, which contrasts with conventional roles of LIF in ES cell pluripotency (Ying et al., 2003). Taken together with SMAD2/3 signaling inhibition, early increases of *Prdm14* observed in iBC generation may be partial to germ cell induction mechanisms. Prdm14 is a transcription factor whose role is both powerful and unclear: reported to be dispensable in BCs yet a major epigenetic regulator expressed in the 2C cleavage stage that may direct lineage commitment in the BC (Yamaji et al., 2008; Luna-Zurita and Bruneau, 2013; Burton et al., 2013). Prdm14 is involved in dynamic biological events that accompany epigenetic reprogramming, such as PGC specification, X chromosome reactivation, and conversion from primed to naive state pluripotency (Yamaji et al., 2008; Gillich et al., 2012; Payer et al., 2013; Kime et al., 2016). Intriguingly, iBC induction with *Prdm14* KD cells proceeded typically for several days but caused iBC-PC cell death at the time when iBC-PCs should begin to polarize and differentiate to iBCs. How Prdm14 is involved in iBC production warrants further investigation.

Notably, our defined conditions resemble germ cell induction medium, suggesting a PGC/germ cell specification process may be involved; the germ line prepares a totipotent genome that is not yet activated epigenetically. Since BCs naturally differentiate from homogenous totipotent cells, the 2C stage ZGA mechanism suggested by MERVL may also play a role (Wu et al., 2017). Supporting this notion, we observed the compacting iBC-PC/iBC originating loci with concomitant *MERVL::*RFP+ cells curiously demonstrating large cleavage-stage cell size and morphology. Some previous studies of MERVL-enriched PSCs touched upon the implication of totipotent hallmarks, ZGA, and 2C-like expression, yet these reports did not demonstrate similar apparent morphological changes (Macfarlan et al., 2012; Blaschke et al., 2013; Ishiuchi et al., 2015). Several reports found ES cell colonies have rare transient MERVL+ cells that return to the ES state. In iBC experiments, we see several cells at iBC-PC loci, or iBC-PC, activating *MERVL::*RFP simultaneously and with relatively consistent sustained expression. We also note that X reactivation suggested by XGFP+ cells among our *MERVL::*RFP+ cells presents a ratio quite different from distinct features of naive PSCs, which should be completely Xa/Xa and have rare MERVL reporter activation (Macfarlan et al., 2012; Payer et al., 2011). Our observation of *MERVL::RFP+* cells with less X reactivation may reflect the varied Xa/Xa state of cleavage stage cells rather than naive PSCs. The enrichment of *Atp1b1* in isolated iBCs was also remarkable since natural embryos activate *Atp1b1* in cleavage stage cells in preparation to act in cell junctions of compaction and as a Na+/K+ ATPase pump subunit to fill the blastocoel (Hamatani et al., 2004; Madan et al., 2007; Stephenson et al., 2010). Isolated iBCs could also show *Zfp42* (Rex1) and *Zscan4*, which are also cleavage stage-induced genes with different roles in pluripotent cells. Moreover, Zfp42 (Rex1) may negatively regulate 2C-related gene expression (Schoorlemmer et al., 2014), yet iBC/iBC-PC-derived ES-like cells have notably increased expression of both Zfp42 (Rex1) and Zscan4 RNAs when compared to ES cells. With these observations, a thorough molecular elucidation of iBC generation would likely improve their quality to obtain full functionality.

Generation of iBCs requires stringent PSC preparation and rounds of iBC purification, yet most of the experiment involves 1 week of simplified defined-media changes. Thus, we envision that iBC technology readily opens avenues in several fields, such as embryology and implantation biology among its promise in early embryogenesis. For instance, even though iBCs cannot develop completely, transplanting iBCs from gene knockdown or knockout PSCs may make it easier to elucidate the molecular mechanisms governing implantation. In any case, our findings may offer a new step in expanding knowledge in pluripotency, totipotency, and embryogenesis, which may be necessary to significantly advancing PSC technologies and related fields.

## AUTHOR CONTRIBUTIONS

Conceptualization, C.K.; Methodology, C.K. and K.T.; Validation, S.O.; Formal Analysis, C.K. and K.T.; Investigation, C.K., K.T., and H.K.; Resources, C.K., K.T., S.Y., and M.T.; Writing - Original Draft, C.K. and K.T.; Writing - Review & Editing, C.K. and K.T.; Visualization, C.K.; Supervision, C.K. and K.T.; Project Administration, C.K.; Funding Acquisition, S.Y., M.A., and M.T.

## ACKNOWLEDGEMENTS

We honor the help of Dr. Hitoshi Niwa for critical input and for providing the Rabbit anti-Mouse CDX2 antibody. We are grateful to Drs. Siqin Bao and Azim Surani for their female XGFP mEpiSC. We also thank Drs. Robert Blelloch and Paul Tesar for providing their mEpiSC for research. MERVL 2C and EOS-S(4+) reporter DNA was provided by Addgene (http://www.addgene.org) under MTA and subcloned into our systems. TROMA-I (Krt8) monoclonal antibody developed by Institut Pasteur was obtained from the Developmental Studies Hybridoma Bank(DSHB), created by the NICHD of the NIH and maintained at The University of Iowa, Department of Biology, Iowa City, IA 52242. We greatly thank the Yamanaka and Takahashi labs for the support and research environment that made this work possible.

## Conflict of Interest

C. K and K.T. have applied for patents related to this technology and extended works. S.Y. is a scientific advisor of iPS Academia Japan without salary.

---

**Figure S1:**
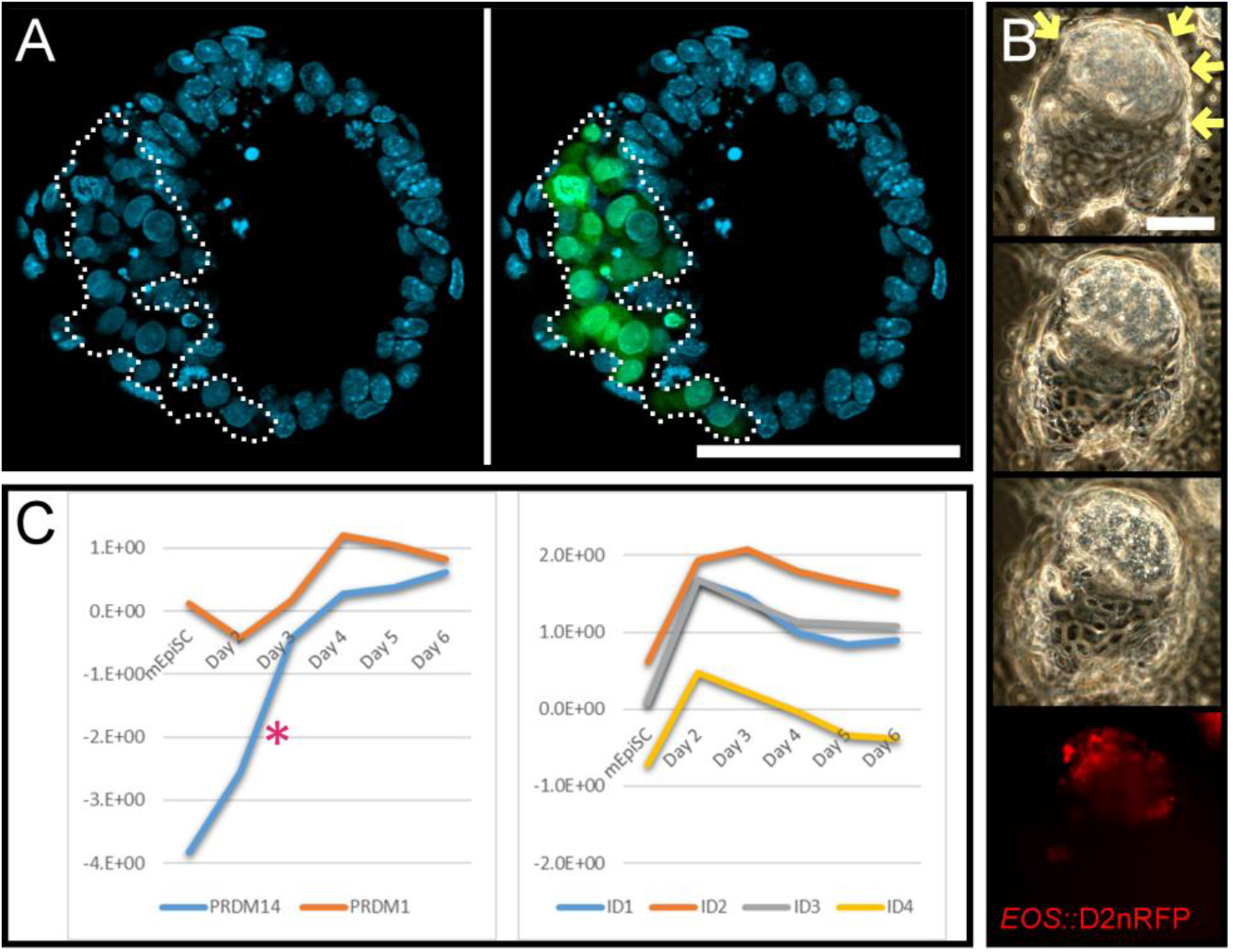
Related to Figure 1 Naïve Conversion Induces Blastocyst-Like Hemispheres and Prdm and Id Gene mRNAs. **A**) Image from Figure 1A structure with XGFP+ cells outlined, +/− the XGFP image layer. XGFP− cells have bright DNA stain punctae indicating condensed DNA in heterochromatin (light blue; Hoechst 33342). XGFP+ cells lack bright DNA punctae. *Scale bar = 100 μm*. **B**) Naïve conversion BC-like hemisphere with *EOS::*D2nRFP+ expression restricted to the putative ICM surrounded by TE-like cells (yellow arrows), across three Z-positions. *Scale bar = 100 μm*. **C**) RT-qPCR of naive conversion time course cDNA samples shows strong induction of *Id* genes in two days (right). *Prdm14* is induced from near undetectable signal in mEpiSC and *Prdm1* is partially maintained before ^~^10-fold increase after 4 days (left). *\*Prdm14 time-course was reported in (Kime et al., 2016)*.

**Figure S2:**
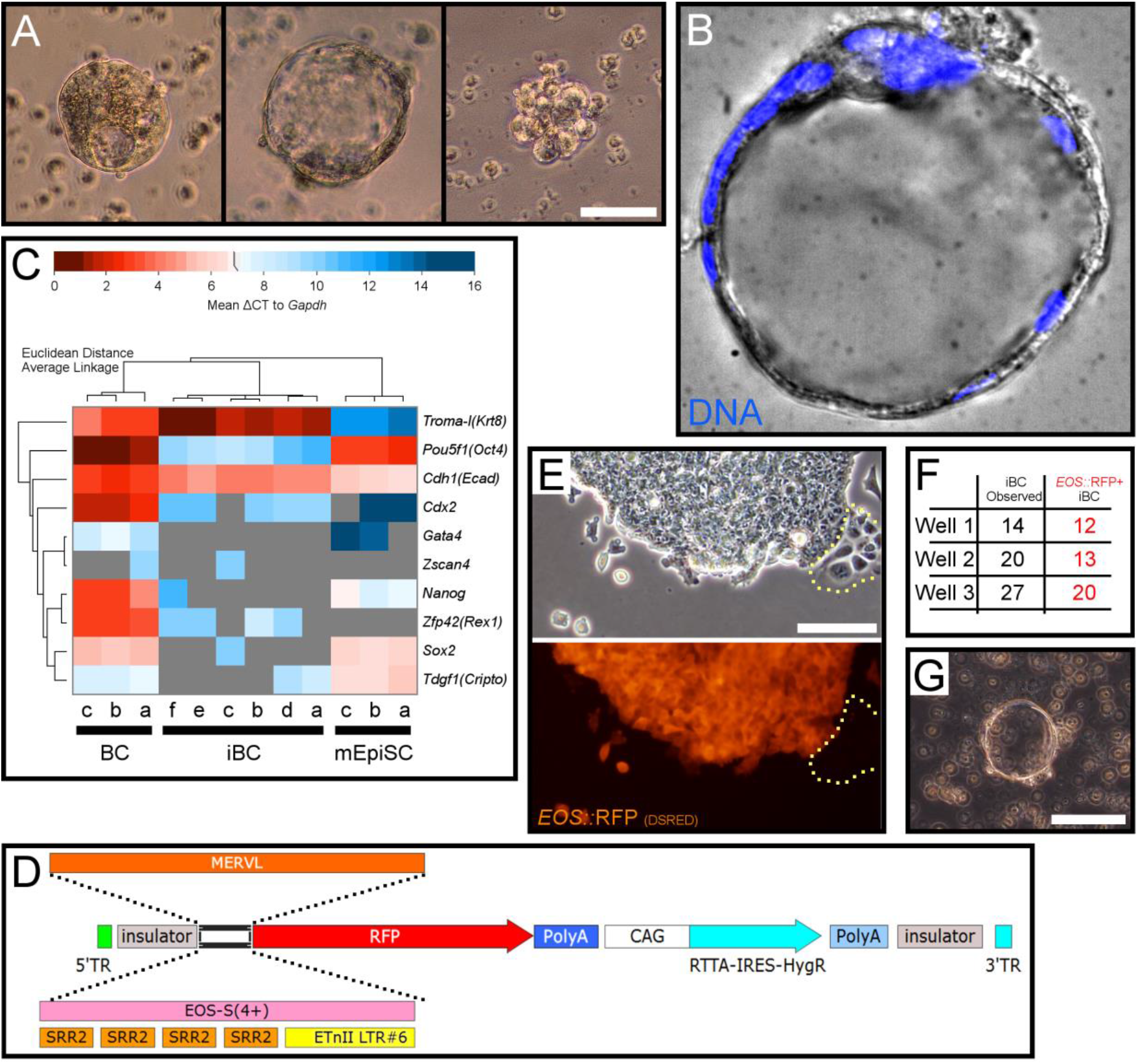
Related to Figure 2, Figure 3, and Figure 7 iBC Characterization and Reporter Constructs. **A**) Early embryo-like structures are released to suspension. *5cale bar (all) = 50 μm*. **B**) Late iBC stained for DNA (Blue; Hoechst 33342). **C**) RT-qPCR of single early BC, early iBC, and mEpiSC cDNA samples, with Euclidean distance and clustering by average linkage, represented as a heat map of ΔCT to *Gapdh*. **D**) Schematic of piggyback reporter systems in this study: *MERVL* or *EOS-S(4+)* synthetic promoters followed by RFP. *RFP used are D5RED, mCherry, or D2nRFP (see methods)*. **E**) *EOS::*RFP+ mEpiSC and *EOS::*RFP− differentiating cells. *Differentiating cells are outlined with yellow dotted line. 5cale bar = 100 μm*. **F**) 6W wells of iBC generation were harvested to ULA plates on Day 6, and counted on Day 7 as total yield and 32 those that were *EOS::*RFP+. *Scale bar = 100 μm*. **G**) iBC induced from another published mEpiSC line (Tesar et al., 2007). *Scale bar = 100 μm*.

**Figure S3:**
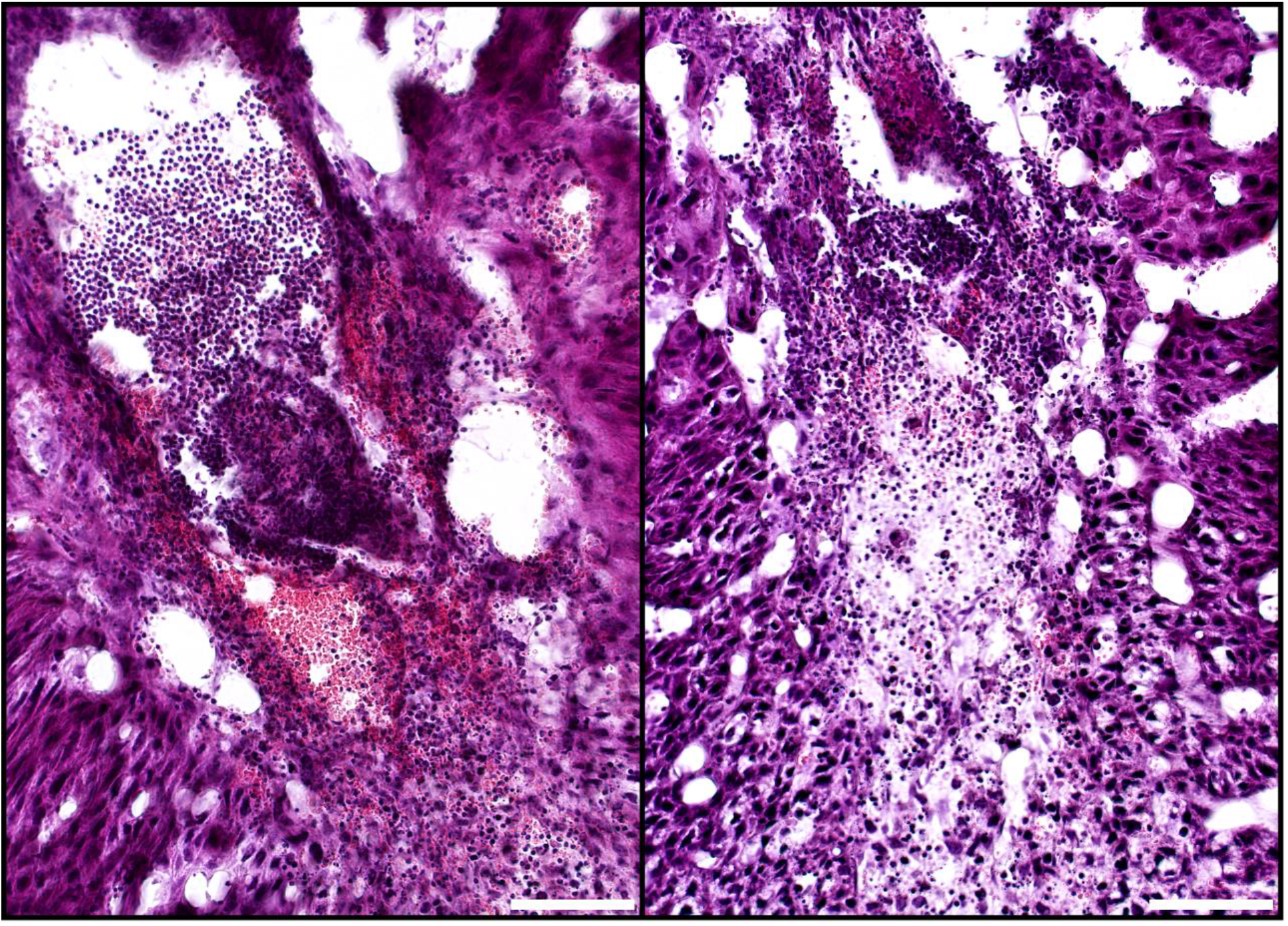
Related to Figure 5 iBCs Implanted Tissues Are Resorbed by Immune Cells. Larger images of H&E stain for E7.5 iBC single source transfer deciduae cryosections. Deciduae had many blood sinuses and ExEm-like cells at the periphery of resorbing tissue. Maternal immune cells were highly present throughout the area, appearing to have accumulated from the blood sinuses. Loosely arranged ExEm-like cells appeared to retract within a degrading embryonic cavity and surrounding small darker stained cells resembling Em cells seen in healthy control embryos (Figure S4). *Scale bars = 100 μm*.

**Figure S4:**
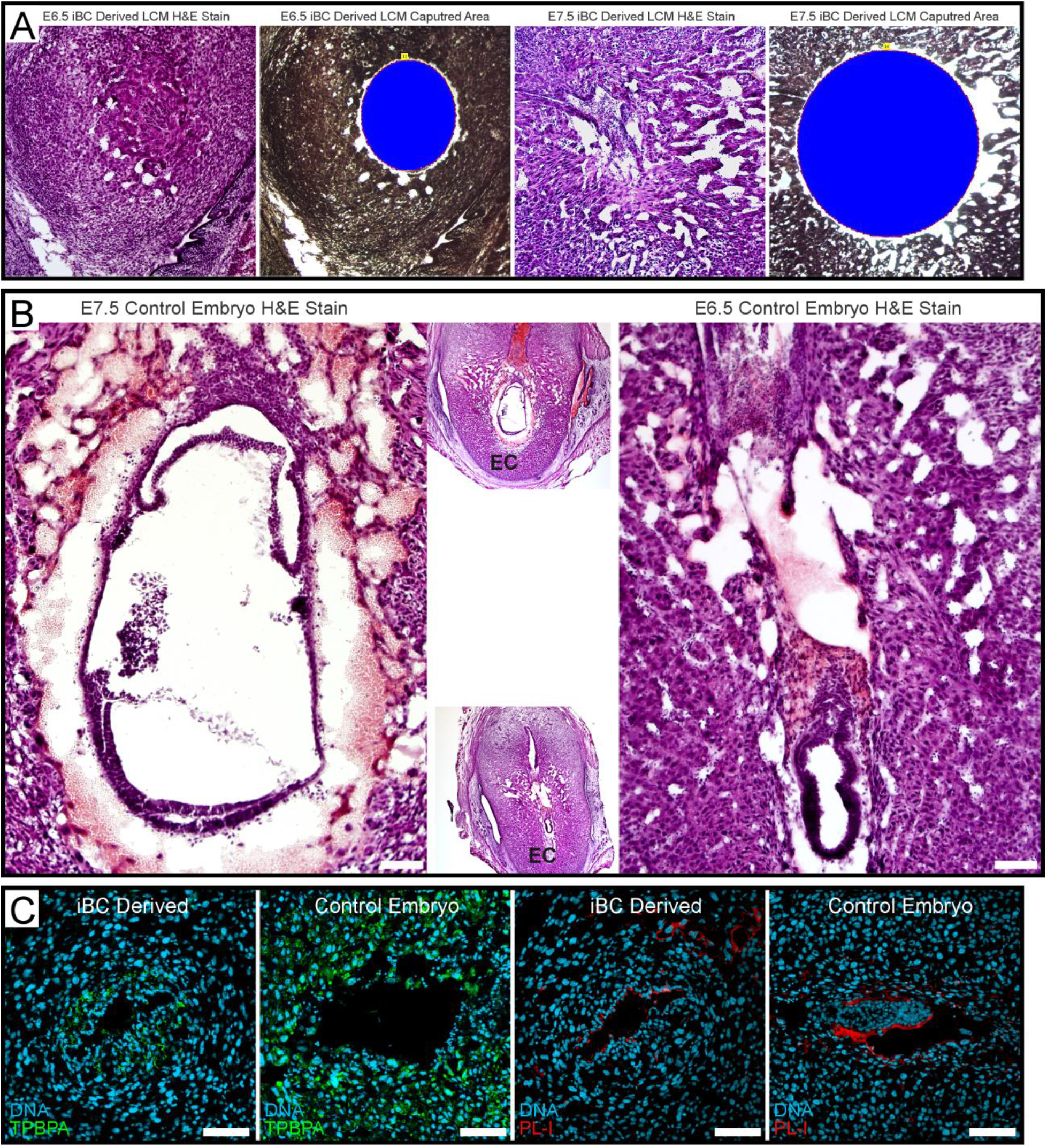
Related to Figure 4, Figure 5. **A**) H&E stain and LCM sampled areas (Blue) of proximal sections from the iBC derived tissue for E7.5 iBC single source uterus transfer from Figure 5A,B, and a E6.5 iBC co-transfer sample. *The LCM samples were used for genomic DNA PCR in Figure 4D*. **B**) H&E stain for E7.5 and E6.5 control embryos for reference. Lower magnification images of the full decidua section with uterine tissue is set to the side. *EC, embryonic cavity. Scale bars = 100 μm*. **C**) Cryosection IHC for post-implantation ExEm lineage markers TPBPA (Green, Left) and PL-I (Red, Right), and DNA (light blue, Hoechst 33342) on E6.5 iBC-derived and control embryo derived deciduae embryonic cavity regions.

**Figure S5:**
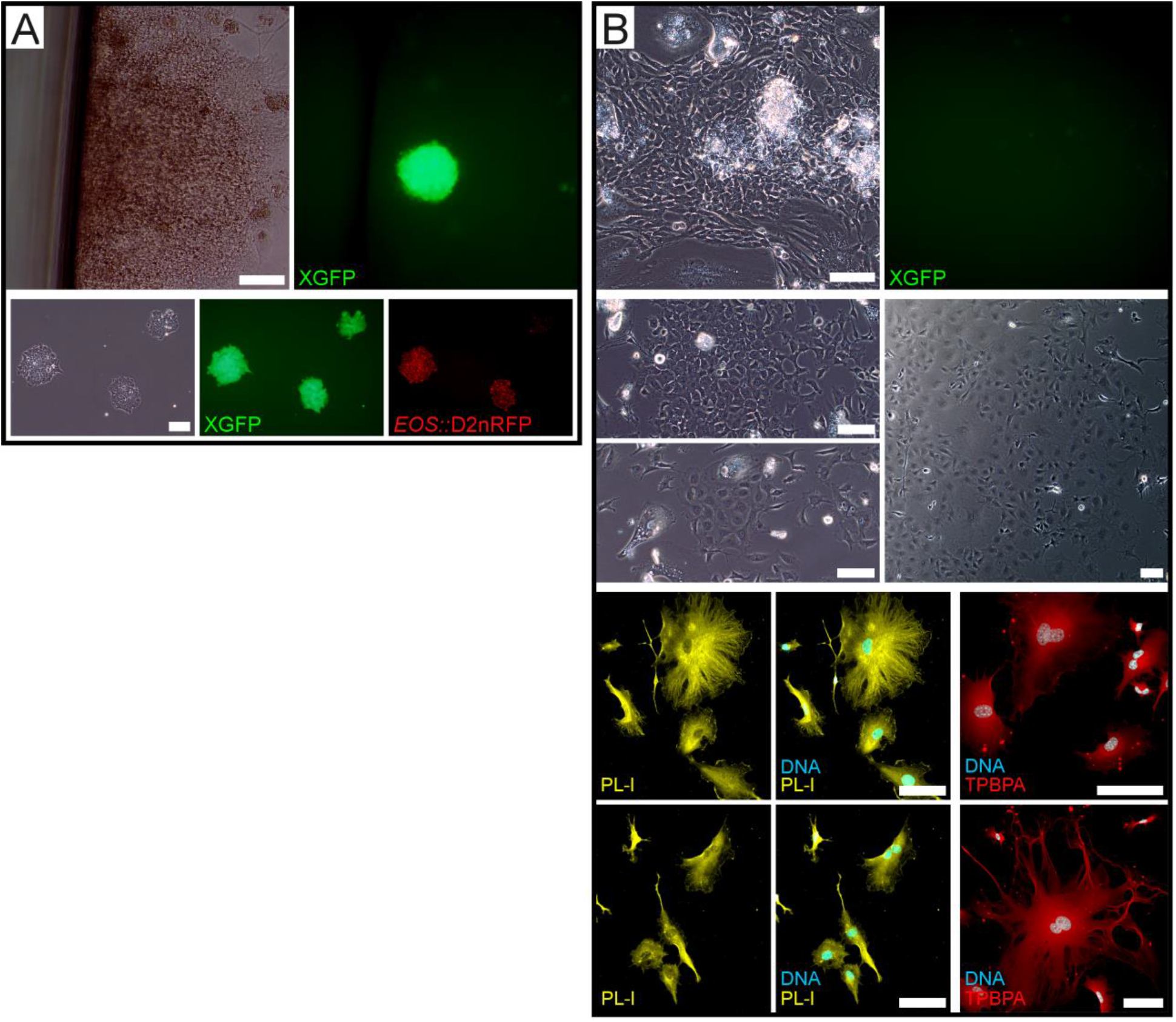
Related to Figure 6 iBC/iBC-PC-Derived Outgrowths Produce ES-Like Cells and TE Lineage Marker-Positive Cells. **A**) *EOS::*D2nRFP mEpiSC induced iBC and iBC-PC were plated on feeders in ES cell derivation conditions. Cells activated the XGFP reporter (upper panel) and stabilized similar to ES cells after several passages and maintained XGFP and EOS::D2nRFP expression. *Scale bars = 100 μm*. **B**) *EOS::*D2nRFP mEpiSC induced iBC and iBC-PC were plated on feeders in TE cell-culture conditions. Outgrowths mostly did not activate the XGFP reporter (upper panel) and could expand for two passages into TE-like and binuclear cells (middle panels). Cells were passaged to slides and stained for post-implantation ExEm cell markers PL-I (yellow, lower left) and TPBPA (red, lower right), and DNA (light blue, Hoechst 33342). *Scale bars = 100 μm*.

**VIDEO S1: Blastocyst-Like Hemisphere Imaged from Z-stack**

A late BC-like hemisphere imaged across the z-dimension, visualized as a composite 3D model and animated for viewing from several angles. XGFP+NANOG+ cells are restricted to a polar mass of the fluid filled dome surrounded by large flat cells with large flat nuclei. Green = XGFP+; Red = NANOG+; Blue = DNA, Hoechst 33342.

**Table S1.** Related to Figure 2 iBC Generation Experiment Outcomes. iBC generation experiments over the course of this study with respect to initial iBC observation and relative outcomes.

| iBC Observed Date * | Relative Yield ** |
| --- | --- |
| 10/25/2014 | + |
| 11/25/2014 | + |
| 12/27/2014 | + |
| 7/14/2015 | + |
| 8/4/2015 | + |
| 10/21/2015 | + |
| 10/28/2015 | + |
| 2/26/2016 | + |
| 3/17/2016 | ++ |
| 3/31/2016 | ++ |
| 5/12/2016 | ++ |
| 6/13/2016 | ++ |
| 9/15/2016 | ++ |
| 10/27/2016 | ++ |
| 11/17/2016 | +++ |
| 1/8/2017 | +++ |
| 2/8/2017 | ++ |
| 4/25/2017 | ++ |
| 5/11/2017 | ++ |
| 6/21/2017 | ++ |
| 12/8/2017 | ++ |
| 12/26/2017 | +++ |
| 1/7/2018 | ++ |

\*May include 1-2 days after because iBC-PC can be harvested each day for approximately 2 days and become iBCs 12-48 hours later in suspension.
\*\*Differences in relative yield reflect experiment optimization changes. Experiments with +++ yield were so abundant that late iBCs readily aggregated and complicated purification.

**Table S2.** Related to Figure 3 iBC vs BC Comparative Microscopy Results. iBC stained for DNA, OCT4, or TROMA-I detection are imaged on LSM700 and LSM800 microscopes (see methods). Three BCs are stained the same and imaged with matching microscope settings to determine detector gains (see methods). EB, mEpiSC Cluster, iBC, and Control Embryo (BC) Single Source and Co-Transfer Uterus Transfer Decidualization Experiments

| LSM700 | Detector Gains | Hoechst | OCT4 | TROMA-I |  | LSM880 | Detector Gains | Hoechst | OCT4 | TROMA-I |
| --- | --- | --- | --- | --- | --- | --- | --- | --- | --- | --- |
| iBC Control Setting |  | 339 | 748 | 406 |  | iBC Control Setting |  | 398 | 731 | 513 |
| Blastocyst-1 |  | 363 | 491 | 540 |  | Blastocyst-1 |  | 356 | 502 | 537 |
| Blastocyst-2 |  | 332 | 447 | 457 |  | Blastocyst-2 |  | 367 | 538 | 585 |
| Blastocyst-3 |  | 375 | 462 | 515 |  | Blastocyst-3 |  | 363 | 601 | 604 |
| LSM700 | Detector Gains | Hoechst | OCT4 | TROMA-I |  | LSM880 | Detector Gains | Hoechst | OCT4 | TROMA-I |
| iBC Control Setting |  | 361 | 552 | 463 |  | iBC Control Setting |  | 293 | 745 | 612 |
| Blastocyst-1 |  | 384 | 414 | 503 |  | Blastocyst-1 |  | 344 | 634 | 630 |
| Blastocyst-2 |  | 403 | 420 | 531 |  | Blastocyst-2 |  | 344 | 609 | 705 |
| Blastocyst-3 |  | 415 | 443 | 557 |  | Blastocyst-3 |  | 354 | 561 | 619 |

**Table S3.** Related to Figure 4 EB, mEpiSC cluster, iBC, and BC Decidualization Experiments. Sterile-male bred pseudopregnant mice (PP2.5) surrogate transfer unit counts and subsequent dissected deciduae counts.

| Control Embryos Only | Control Embryos | n/a | Deciduae Observed | # above control | Control Decidualization Frequency |
| --- | --- | --- | --- | --- | --- |
|  | 7 | 0 | 6 | 0 | 86% |
|  | 7 | 0 | 0 | 0 | 0% |
|  | 7 | 0 | 7 | 0 | 100% |
|  | 7 | 0 | 3 | 0 | 43% |
|  | 10 | 0 | 10 | 0 | 100% |
|  | 10 | 0 | 10 | 0 | 100% |
|  | 4 | 0 | 0 | 0 | 0% |
| iBCs Only | Control Embryos | iBCs | Deciduae Observed | # above control | iBC Decidualization Frequency |
|  | 0 | 10 | 0 | 0 | 0% |
|  | 0 | 10 | 6 | 6 | 60% |
|  | 0 | 10 | 0 | 0 | 0% |
|  | 0 | 10 | 0 | 0 | 0% |
|  | 0 | 10 | 1 | 1 | 10% |
|  | 0 | 8 | 0 | 0 | 0% |
|  | 0 | 10 | 2 | 2 | 20% |
|  | 0 | 11 | 0 | 0 | 0% |
|  | 0 | 10 | 0 | 0 | 0% |
|  | 0 | 10 | 0 | 0 | 0% |
|  | 0 | 10 | 0 | 0 | 0% |
|  | 0 | 10 | 1 | 1 | 10% |
|  | 0 | 10 | 0 | 0 | 0% |
|  | 0 | 10 | 0 | 0 | 0% |
|  | 0 | 10 | 0 | 0 | 0% |
| iBC + Control Co-Transfer | Control Embryos | iBCs | Deciduae Observed | # above control | Control Decidualization Frequency * |
|  | 4 | 8 | 5 | 1 | 125% |
|  | 5 | 5 | 7 | 2 | 140% |
|  | 5 | 5 | 0 | 0 | 0% |
|  | 5 | 5 | 4 | 0 | 80% |
|  | 4 | 7 | 5 | 1 | 125% |
|  | 5 | 5 | 8 | 3 | 160% |
|  | 5 | 5 | 6 | 1 | 120% |
|  | 5 | 5 | 5 | 0 | 100% |
|  | 5 | 6 | 4 | 0 | 80% |
|  | 5 | 6 | 6 | 1 | 120% |
|  | 4 | 8 | 4 | 0 | 100% |
|  | 4 | 9 | 0 | 0 | 0% |
|  | 4 | 10 | 5 | 1 | 125% |
|  | 4 | 8 | 6 | 2 | 150% |
|  | 4 | 8 | 5 | 1 | 125% |
|  | 4 | 8 | 5 | 1 | 125% |
|  | 4 | 8 | 5 | 1 | 125% |
|  | 3 | 7 | 4 | 1 | 133% |
|  | 2 | 8 | 0 | 0 | 0% |
|  | 4 | 8 | 7 | 3 | 175% |
|  | 4 | 7 | 3 | 0 | 75% |
|  | 4 | 8 | 8 | 4 | 200% |
|  | 4 | 8 | 3 | 0 | 75% |
|  | 4 | 8 | 3 | 0 | 75% |
|  | 4 | 8 | 5 | 1 | 125% |
|  | 4 | 8 | 3 | 0 | 75% |
| **EB + Control Co-Transfer** | **Control Embryos** | **EB** | **Deciduae Observed** | **# above control** | **Control Decidualization Frequency \*** |
|  | 4 | 8 | 0 | 0 | 0% |
|  | 4 | 7 | 0 | 0 | 0% |
| **EB Only** | **Control Embryos** | **EB** | **Deciduae Observed** | **# above control** | **EB Decidualization Frequency** |
|  | 0 | 10 | 0 | 0 | 0% |
|  | 0 | 7 | 0 | 0 | 0% |
|  | 0 | 10 | 0 | 0 | 0% |
|  | 0 | 10 | 0 | 0 | 0% |
| **mEpiSC Cluster + Control Co-Transfer** | **Control Embryos** | **mEpiSC Clusters** | **Deciduae Observed** | **# above control** | **Control Decidualization Frequency \*** |
|  | 4 | 8 | 2 | 0 | 50% |
|  | 4 | 8 | 3 | 0 | 75% |
|  | 4 | 8 | 4 | 0 | 100% |
|  | 4 | 8 | 2 | 0 | 50% |
| **mEpiSC Cluster Only** | **Control Embryos** | **mEpiSC Clusters** | **Deciduae Observed** | **# above control** | **mEpiSC Decidualization Frequency** |
|  | 0 | 10 | 0 | 0 | 0% |
|  | 0 | 10 | 0 | 0 | 0% |
|  | 0 | 10 | 0 | 0 | 0% |
|  | 0 | 10 | 0 | 0 | 0% |

\*Control Embryos are the positive control value. Calculations assume control embryo deciduae and maximal 100% result. Deciduae in excess of 100% obviate iBC contribution.

## METHODS

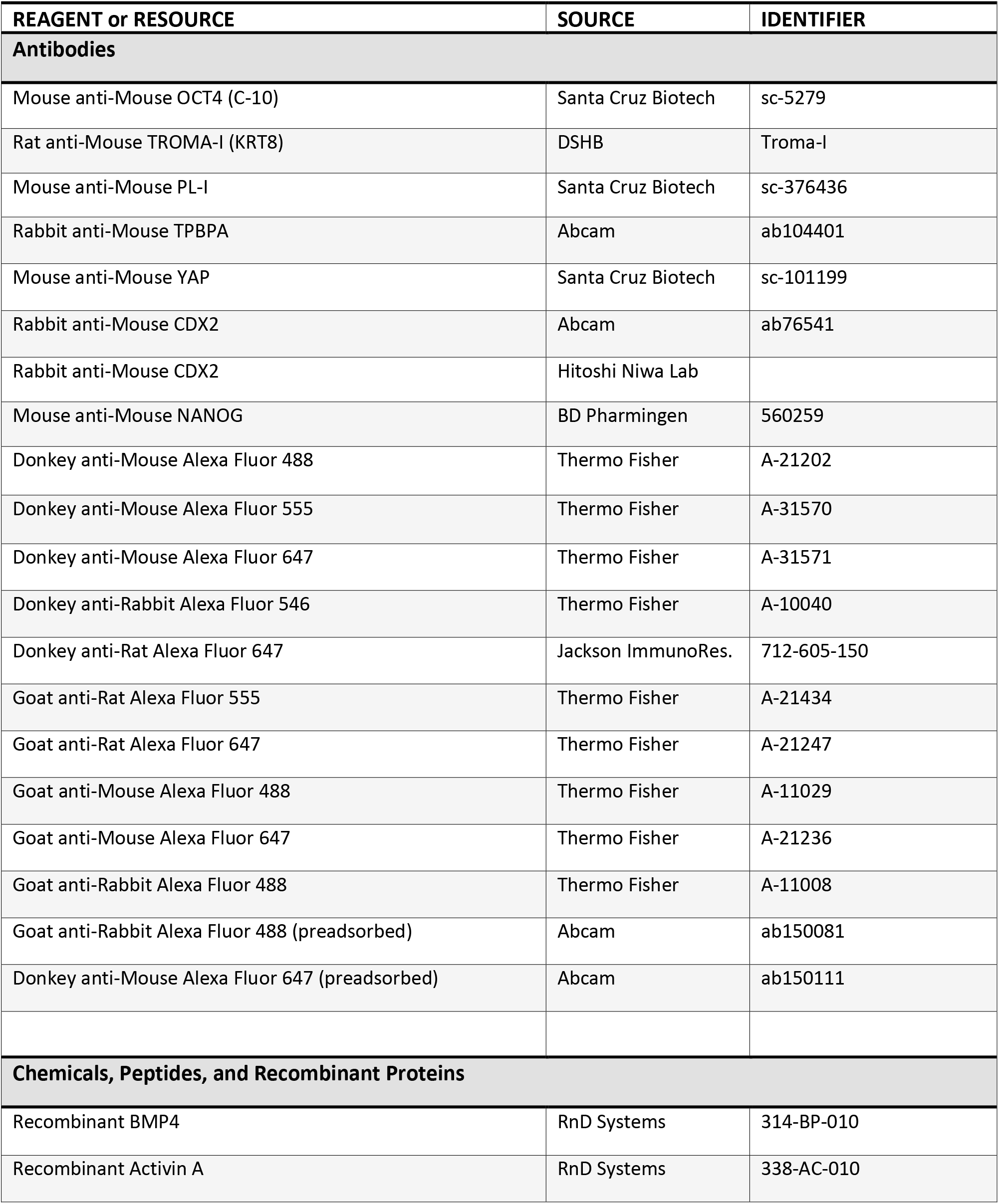
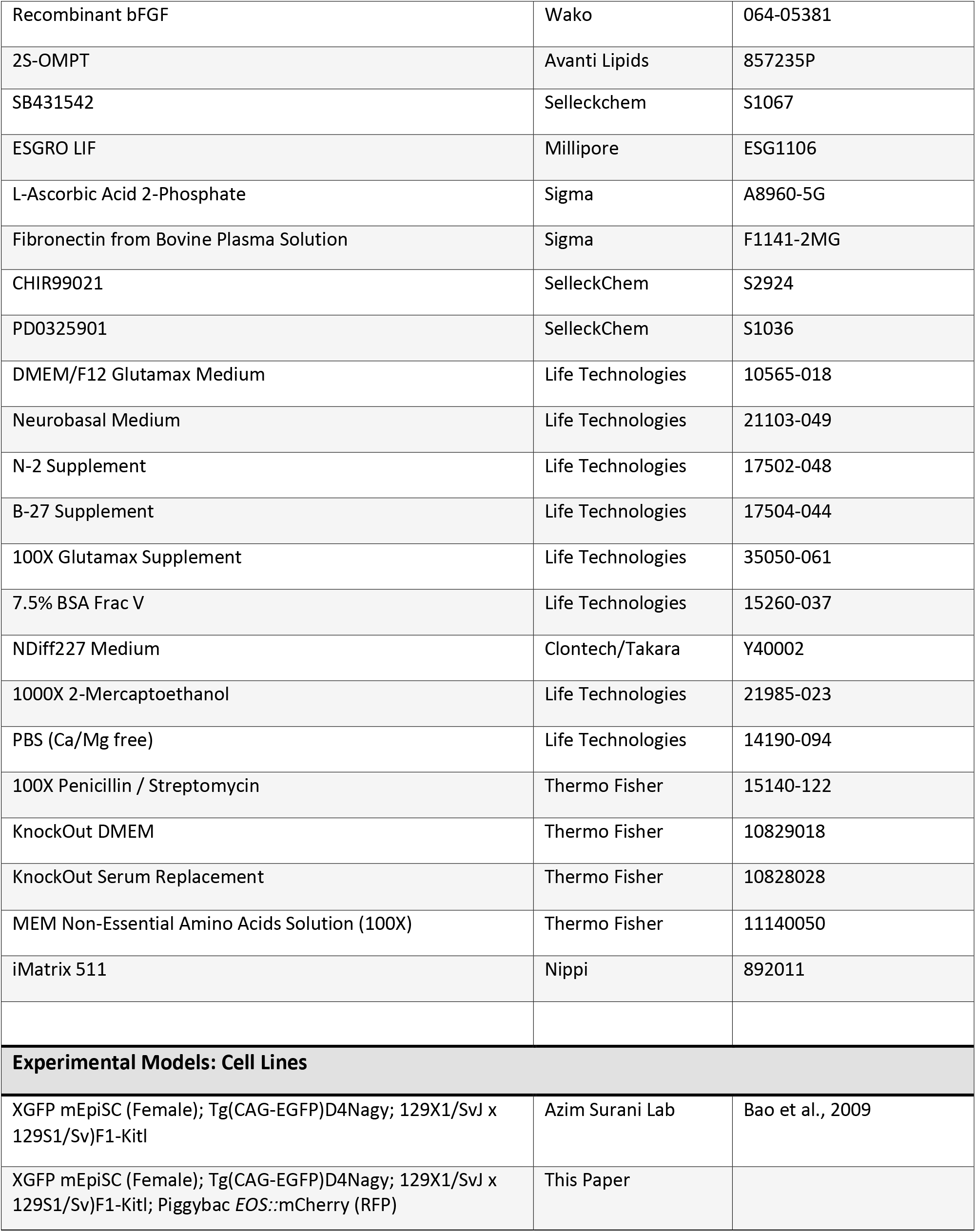
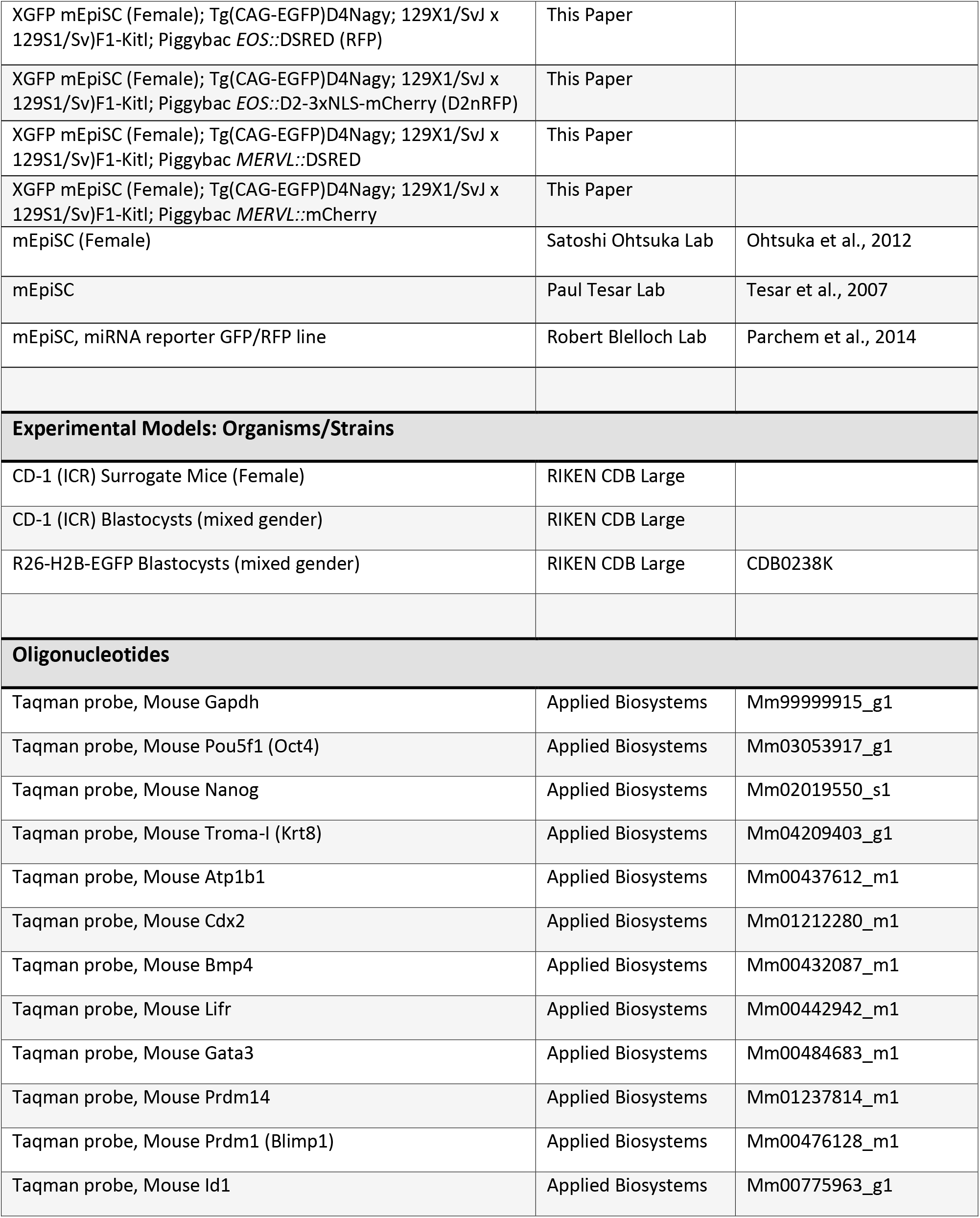
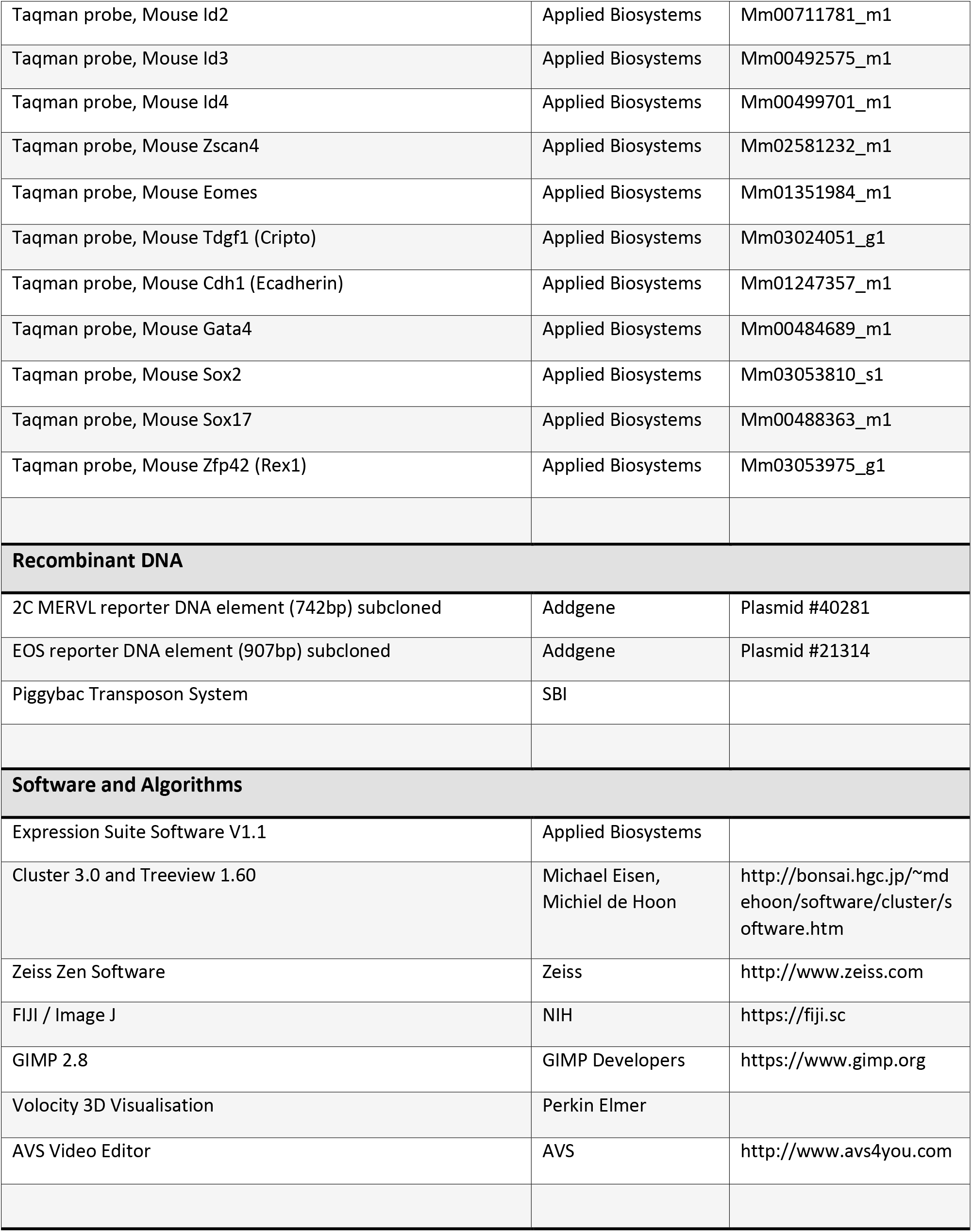
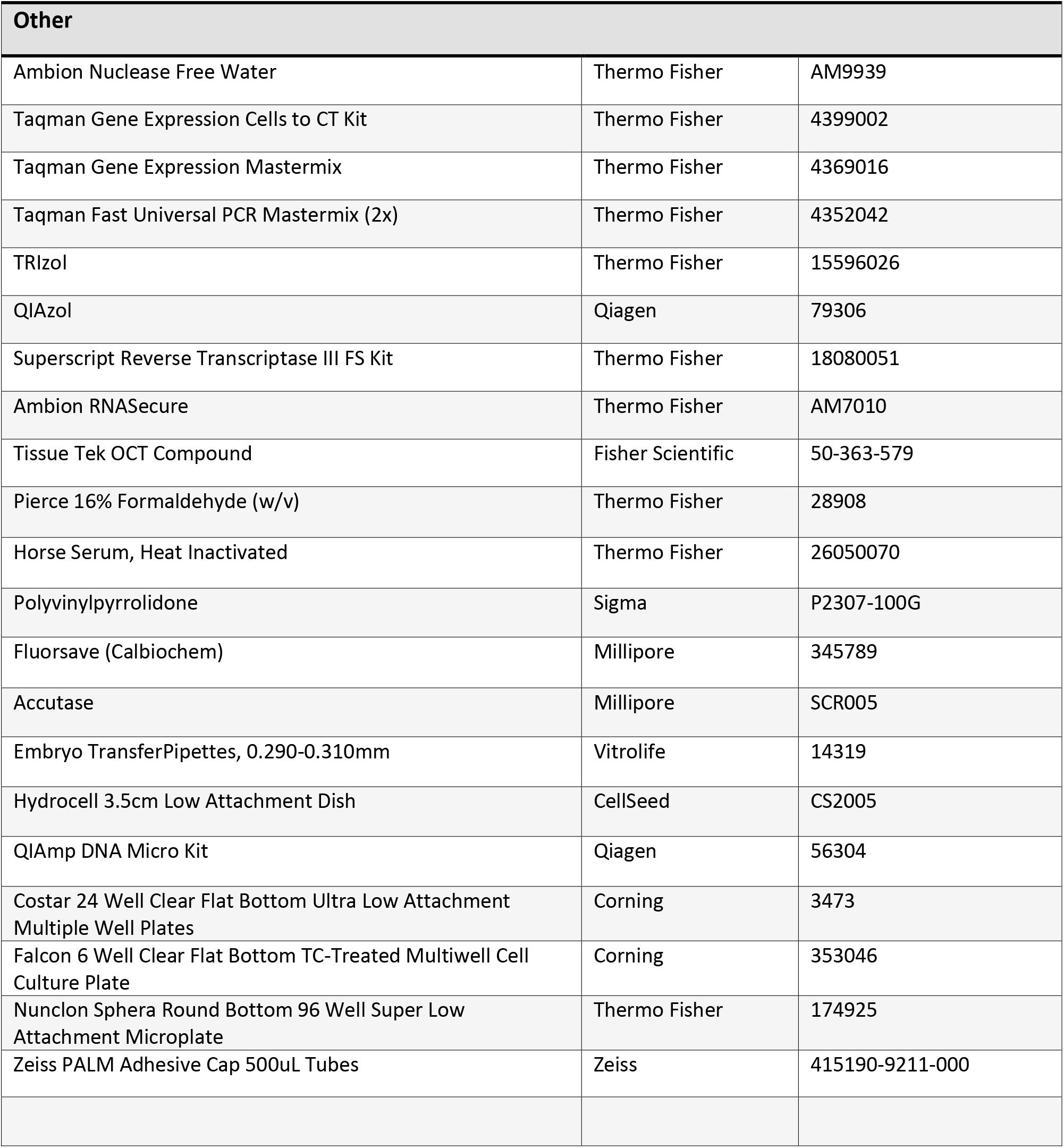

### Contact for Reagent and Resource Sharing

Further information and requests for reagents can be directed to Cody Kime. MTAs required.

### Experimental Model Details

#### Animal Use

Mouse handling and experiments were carried out with humane methods approved by RIKEN Kobe Safety Center. Sterile-male bred pseudopregnant surrogate CD-1 (ICR) female mice were prepared at PP2.5 and then control BCs, iBCs, mEpiSC clusters, or EBs were transferred to the uterus using standard embryo IVF pipetting techniques. CD-1 (ICR) BCs and R26-H2B-EGFP BCs were used for control BC experiments (Abe et al., 2011).

#### mEpiSC Culture

mEpiSC Culture Media (MCM): NDiff227 supplemented with 20 ng/mL ActivinA, 12 ng/mL bFGF, and 1:100 penicillin/streptomycin. Media and supplements were stored separately at −20 °C in aliquots and thawed fresh at least every 4 days and stored at 4 °C. mEpiSC were cultured on plates coated for 1 hour at room temperature with 1:100 Fibronectin:PBS. Medium was changed daily, and cells were passaged as small clumps every 2-3 days at ^~^1:10-20, never exceeding 30% confluent. Cell colonies remained less than 200-300 μm wide and largely resembled homogenous mEpiSC colonies with few single cells. Cell passage was carried out, in brief, with PBS wash, fresh Accutase for 55 seconds, PBS wash, 2 mL of MCM, scraped, triturated 6-8 times in a conical vial, then dispersed ^~^1:10-20 in MCM. If cells exhibited signs of differentiation the culture was discarded and replaced by a freshly thawed stock.

## Method Details

### CTSFES Media Preparation for Working Media

*CTSFES Basal Media ^~^1L Preparation:* [500 mL DMEM:F12+Glutamax, 500 mL Neurobasal Media, 10 mL B27 Supplement, 5 mL N2 Supplement, 5 mL Glutamax Supplement, 670 μL 7.5% BSA Frac V Solution]; filtered at 0.22 μm, aliquoted, and stored immediately at −20 °C; thawed overnight at 4 °C and used for 1-8 days.

*CTSFES ‘Working Medium’ after thawing for experimental use:* Add 1:100 penicillin/streptomycin, 1:1000 2-ME, 64 μg/mL ascorbic acid 2-phosphate.

### mEpiSC Preparation for Naive Conversion or iBC Generation

Target wells of 6W plate were coated with 1.5 mL 1:100 Fibronectin:PBS substrate for 1 hour at room temperature. Stock cultures of near-passage mEpiSC colonies were sourced for passage into conversion experiment as follows: PBS wash, freshly thawed room temperature Accutase for 1 minute, Accutase gently aspirated, washed again with equal volume of PBS while tapping the plate to release single-cells, PBS wash gently aspirated, 37 °C prewarmed fresh Accutase was added and incubated at 37 °C for 5-7 minutes until cells floated and dispersed freely. 5X volume 1:1 PBS:MCM was added and the volume triturated 10-20 times in 15 mL conical vial. Cells were centrifuged at 200xG for 3 minutes. mEpiSC pellet was resuspend in 1-2 mL of MCM and live cells were counted. Cells were diluted in MCM to yield ^~^20,000 cells/1.5 mL for naive conversions or 30-50,000 cells/1.5 mL for iBC generation, mixed evenly. Fibronectin:PBS coating was aspirated from target plates and 1.5 mL of diluted cells in MCM were added per well. Cells were incubated at 37 °C for 14-16 hours before conversion media is added; plates were often checked 2-3 hours after plating to ensure cells plated as single evenly dispersed cells.

### Naive Conversion Experiment

*Naive Conversion Experiment Media(NCM) (8 days of changes):* Working Medium + [10 ng/mL BMP4, 1000 units/mL ESGRO LIF, and 1 μM OMPT; prepared fresh at least every 4 days.

6W wells plated with ^~^20,000 mEpiSC cells/well were fed 2 mL of NCM daily starting ^~^14-16 hours after cells were plated with the preparation noted prior in these methods.

### Generation Experiment

*iBC Generation Media Phase 1, Day 0-3 Media (4 changes):* Working Medium + [10 ng/mL BMP4 and 1 μM SB43152]; prepared fresh on Day 0, and SB431542 is increased to 3uM for Days1-3. *iBC Generation Media Phase 2, Day 4-6 Media (3 changes):* Working Medium + [5 ng/mL BMP4, + 500-1000 units/mL ESGRO LIF, and 0.5-1 μM OMPT]; prepared fresh on Day 4.

6W wells plated with 30-50,000 mEpiSC cells/well were fed 2 mL of Phase 1 Medium daily at a similar time, starting 14-16 hours after cells were plated with the preparation noted prior in these methods. From Day 4, 2 mL of Phase 2 medium was changed daily. On Day 6 and Day 7, iBC-PCs and some emerging iBCs were collected with ART P1000G Wide Bore Pipette tips. iBC Generation Plate was leaned at a 45° angle, and the upper 1 mL (primary) was harvested to one well of a 24-well ULA Plate; the lower 1 mL (secondary) was drawn up and cascaded over the plate once and then harvested to a separate well of a 24-well ULA Plate. 2 mL of Phase 2 medium was replaced on the plate if the culture was observed or used later. *Some iBC experiments included 0.2 μM sodium pyruvate. In a few experiments, 5B431542 was varied between 1 and 10 μM, and Phase 1 and Phase 2 media were mixed 1:1 on Days 3, 4, or 5*.

Primary and secondary harvests from one 6W well of iBC generation were considered together, although secondary harvests contained more iBC-PCs, iBCs, and cell debris. Early on Day 7, primary and secondary harvests were observed for brief periods, and the emergence of morula-like structures and early blastocyst-like structures from iBC-PCs was noted on the 24W ULA plate lid. Working medium or Phase 2 medium was placed in a Hydrocell 3.5-cm plate and incubated for 1 hour at 37 °C. iBCs were judged by morphology for blastocyst-like characteristics and isolated by embryo transfer pipette to the Hydrocell 3.5-cm plate and incubated for 1-3 hours at 37 °C. The Hydrocell 3.5-cm plate of near-completely purified iBCs were then sourced for analysis or IVF transfer into PP2.5 sterile-male bred pseudopregnant mice. When iBCs were transferred to pseudopregnant mice, they were washed 3 times by transfer into separate drops of standard embryo transfer medium. In such transfers, unique glass pipettes were used between each step to ensure sample handling was accurate.

### iBC/iBC-PC Outgrowth Experiments

6-well plates were coated with iMatrix511 and then plated with feeder cells and incubated overnight. The feeders were evenly plated and freshly prepared 2iLIF or CDM-FAXY was changed in at 1 mL/well from medias prepared as follows:

**2iLIF:** 200 mL CTSFES Basal Media + additional 1.2 mL 7.5% BSA Frac V Solution, 1000 units/mL ESGRO LIF, 3 μM CHIR99021, and 1 μM PD0325901, prepared fresh every 4 days.
**CDM-FAXY:** 200 mL of CTSFES Basal Media + 1.2 mL 7.5% BSA Frac V Solution, with supplements as published previously (Ohinata and Tsukiyama, 2014) except with 1:1000 2-ME in place of monthioglycerol.

iBC and iBC-PC were purified by pipette and combined. The combined structures were pipetted against the bottom of the tube to break them up and then plated in the wells of 2iLIF or CDM-FAXY media, changed every other day. After one week, plates were replated in their respective medias on fresh feeders on 6-well plates coated with iMatrix511. 2iLIF cultures were then fed media daily with cell passage thereafter on iMatrix511 coated plates without feeders. CDM-FAXY cultures were fed every other day, replated once more onto iMatrix511 coated plates without feeders and then twice thereafter on fibronectin coated plates.

### Embryoid Body Formation Experiment

*EB Medium ^~^100mL:* [80 mL KnockOut DMEM, 20 mL KnockOut Serum Replacement, 1:1000 2-ME, 1 mL NonEssential Amino Acids Solution, 1 mL Glutamax, 1:100 penicillin/streptomycin]; prepared fresh.

mEpiSC were prepared as single cells as in preparation for Naive Conversion or iBC Generation until pelleted. The pellet was resuspended in 1 mL of EB Medium, and live cells were counted. Cells were plated at 100 μL/well in 96-Well Nunclon Sphera Super Low Attachment Microplates, at 500, 1000, or 2000 cells/well and incubated at 37 °C. On Day 4, 100 μL of additional EB medium was added per well. EBs were collected on Day 8, and smaller sized and evenly formed EBs were sourced for IVF transfer into PP2.5 sterile-male bred pseudopregnant mice. When EBs were transferred to pseudopregnant mice, they were washed 3 times by transfer into separate drops of standard embryo-transfer medium.

### Mouse Decidua Dissection and Cryosectioning

Surrogate mice were sacrificed humanely by standard protocol at E5.5-E9.5, as estimated by transfer timing at PP2.5. Mice were viewed from ventral side and the abdominal area was dissected to present the uterus horns to the fore and posterior of the mouse. Distinct deciduae were counted and noted on mouse cards and dissections were imaged with Sony Xperia 3 S0-01G. When deciduae were desired for cryosection, they were dissected from uterine tissue to individual deciduae, washed in DPBS, and then fixed overnight in paraformaldehyde at 4 °C. Fixed deciduae were washed with PBS and then gradually desiccated with 30% Sucrose/PBS solution overnight at 4 °C and then washed and placed in OCT Compound. Deciduae from one uterus horn were pooled into one sectioning mold, labeled, and stored at −80 °C in OCT compound. OCT compound molds of deciduae were later placed in the Microm HM560 microtome at −20 °C and sections are cut at 10-30 μM thickness and placed on slide glass, dried for 1-2 hours under room temperature blown air, dried 1 hour at 37 °C, then stored at −30 °C in sealed slide containers until later use. Figure S4C sections were cut at 5 μM thickness.

### Hematoxylin and Eosin (H&E) Staining

Previously prepared cryosections of deciduae were thawed from frozen slides and rinsed in PBS, stained with hematoxylin for 5 minutes, eosin for 2 minutes, and then washed with increasing mixed alcohol concentrations and then xylene before finalization in malinol with coverslips sealed by nail polish.

### Fluorescence Imaging and Confocal Microscopy

Immunocytochemistry in Figure 1 A,B, and Figure S1A was carried out by fixation with paraformaldehyde and then blocking and staining in 5% BSA/PBS with mouse anti-mouse Nanog 1:200; then donkey anti-mouse Alexa Fluor 555 1:200, followed by Hoechst 33342 1:1000; imaged in PBS with Zeiss LSM 510 Confocal Microscope. Z-stack images from these samples were used for Supplemental Video 1 by visualization in Velocity software, exported to video and labeled with AVS Video Editor which processed the 16:9 aspect ratio and reduced data size. Figure 1C was prepared using the same methods with 3 μg/mL of rat anti-mouse TROMA-I and mouse anti-mouse Nanog 1:200; then goat anti-rat Alexa Fluor 555 1:500 and donkey anti-mouse Alexa Fluor 647 (Thermo) 1:500, followed by Hoechst 33342 1:1000.

Live-cell XGFP fluorescence was imaged in Figure 1D Zeiss Z1 microscope.

DNA stain and imaging in Figure S2B were carried out by fixation with paraformaldehyde, permeabilization in 0.2% Triton X-100, and blocking and staining in 2% FBS/PBS; Hoechst 33342 1:2000, and imaged in 20% glycerol/PBS suspension slide with Olympus Confocal Microscope (CSU-X1).

Immunocytochemistry in Figure 2D,E, Figure 3B,C was carried out by washing BCs or iBCs or iBC-PCs in 3 mg/mL polyvinylpyrrolidone in PBS, fixation with paraformaldehyde, permeabilization in 0.25% Triton X-100, and blocking and staining in 4% horse serum/PBS. Figure 2D,E samples were stained with 1:100 mouse anti-mouse YAP; then 1:500 goat anti-mouse Alexa Fluor 647. Figure 3B samples were stained 1:100 mouse anti-mouse YAP and 1:100 rabbit anti-mouse CDX2(Abcam); then 1:500 goat anti-mouse Alexa Fluor 647 and 1:500 goat anti-rabbit Alexa Fluor 488 (Thermo). Figure 3C samples were stained with 3.6 μg/mL rat anti-mouse TROMA-I and mouse antimouse OCT4 1:200; then goat anti-mouse Alexa Fluor 488 1:500 and goat anti-rat Alexa Fluor 647 1:500. All samples were followed by Hoechst 33342 stain 1:1000-2000 and imaged in 20% glycerol/PBS suspension slide with Zeiss LSM 700 or 880 Confocal Microscopes. Samples were washed with blocking buffer ^~^3 times between fixing and staining stages.

IHC in Figure 5B was carried out by thawing previously cryosectioned deciduae slides, washing with PBS, permeabilization with 0.2% Triton X-100, and then blocking and staining in either 2% BSA/PBS or 4% horse serum/PBS with 1.8-9 μg/mL rat anti-mouse TROMA-I; then donkey anti-rat Alexa Fluor 647 1:250-500, followed by Hoechst 33342 1:2000, sealed in Fluorsave and imaged with Zeiss LSM 700 Confocal Microscope.

Immunocytochemistry in Figure 6C was prepared by fixation with paraformaldehyde, permeabilization in 0.25% Triton X-100, and blocking and staining in 4% horse serum/PBS. Antibodies were used in separate stains:

1:100 mouse anti-mouse OCT4; then 1:500 goat anti-mouse Alexa Fluor 647.
1:100 mouse anti-mouse NANOG, then 1:500 goat anti-mouse Alexa Fluor 647.
1:100 mouse anti-mouse YAP, then 1:500 goat anti-mouse Alexa Fluor 647.

All staining was followed by Hoechst 33342 1:2000 and imaged with Zeiss LSM 700 Confocal Microscope. Samples were washed with blocking buffer ^~^3 times between fixing and staining stages.

Immunocytochemistry of TE-like cells in Figure 6C was prepared by fixation with paraformaldehyde, permeabilization in 0.2% Triton X-100, and blocking and staining in 4% horse serum/PBS with rabbit anti-mouse CDX2 (gift from Hitoshi Niwa lab) 1:1000; then goat anti-rabbit Alexa Fluor 488 (Thermo) 1:500, followed by Hoechst 33342 1:2000 and imaged in PBS with Zeiss LSM 700 Confocal Microscope. Sample was washed with blocking buffer ^~^3 times between fixing and staining stages.

IHC in Figure S4C was carried out by thawing previously cryosectioned deciduae slides, air dried, washing with PBS, permeabilization with 0.1% Triton X-100, and then blocking in 2.5% skim milk for 30 min. The blocked sections were stained separately as follows:

1:100 mouse anti-mouse PL-I; then 1:1000 donkey anti-mouse Alexa Fluor 647 (Abcam).
1:100 rabbit anti-mouse TPBPA; then 1:1000 goat anti-rabbit Alexa Fluor 488 (Abcam).

All staining was followed by DAPI for nuclear DNA visualization and imaged on both the LSM 700 Confocal Microscope and Keyence BZ-X700. Samples were washed with blocking buffer ^~^3 times between fixing and staining stages.

Live cell RFP and GFP fluorescence was imaged in Figure 3D, Figure 7D,E,F,G, Figure S1B, Figure S2E, and Figure S5A,B, with a Olympus IX71 Microscope.

Immunocytochemistry in Figure S5B was prepared by fixation with paraformaldehyde, permeabilization in 0.25% Triton X-100, and blocking and staining in 4% horse serum/PBS. Antibodies were used in separate stains:

1:100 mouse anti-mouse PL-I; then 1:500 donkey anti-mouse Alexa Fluor 488.
1:100 rabbit anti-mouse TPBPA, then 1:500 donkey anti-rabbit Alexa Fluor 546.

All staining was followed by Hoechst 33342 1:2000 and imaged with Zeiss LSM 700 Confocal Microscope. Samples were washed with blocking buffer ^~^3 times between fixing and staining stages.

### Comparative Microscopy

Zeiss LSM700 and LSM880 confocal microscopes were used as indicated in Table S2. For each iBC sample, channel laser intensities, pinhole, and objective were selected to produce a clear image. The channel_gains were determined manually by setting each channel with range indicators and increasing gain until few target pixels saturated the signal, and then the image was captured. For comparison, three BCs prepared with identical methods were imaged with the same microscope, laser intensity, pinhole, and objective settings as the compared iBC image. The channel gains were determined manually by the same method of using the range indicator setting and increasing the gain until few target pixels saturated the signal, and then the image was captured.

### Light Microscopy

Bright field and phase contrast microscopy were carried out on several microscope models. When accompanied by or prepared as composite in fluorescent images, the same microscope was used. For all others, imaging was carried out as follows: Figure S2A images were taken with Olympus CKX41 Microscope. Figure 2B,C, Figure 3D, Figure 6C (left panel), Figure 7B,D,E,F,G, Figure S1B, Figure S2E,G, Figure S5A,B, were taken with Olympus IX71 Microscope.

H&E Stained cryosection slides in Figure 5A,C, Figure S3, Figure S4A,B, were imaged with Olympus IX71 Microscope

Sony Xperia 3 S0-01G was used for Figure 4B.

Zeiss Laser Palm Microbeam was used for imaging Figure S4A(right panels).

### Recombinant DNA Preparation

We prepared RFP as DSRED, mCherry, and also a modified mCherry under the *EOS* reporter by adding a mouse ornithine decarboxylase destabilization domain(D2) and nuclear localization tags to the RFP(*EOS::*D2nRFP; Li et al., 1998). D2 drastically reduces the half-life of the D2nRFP, providing timely live RFP responsiveness to mRNA level changes: the D2nRFP signal more closely represents Oct4/Sox2 heterodimer transcriptional activity. We also cloned RFP under the 2C *MERVL* reporter promoter. All reporter systems were cloned in piggybac vector systems with 5’ and 3’ insulators.

### Laser Capture Microdissection and gDNA PCR

Unique primers for hygromycin resistance transgene were designed using NCBI Primer Blast web software and optimal primers were selected. Jackson Labs (JAX) universal mouse genomic DNA primers were also used for control PCR. H&E stained cryosection samples of interest were prepared using standard slide cover removal techniques and then automated LCM with the Zeiss PALM Microbeam with close cut parameters. Selected tissues were collected with adhesive cap, 500 μl tubes. Genomic DNA was purified from collected tissues using the Zeiss PALM Protocols DNA Handling manual page 21 with a QIAmp DNA Micro Kit, eluting in 20ul of nuclease free water. 5 μl of purified sample was used in 25 μl PCR reactions with touchdown thermocycling using the following DNA oligonucleotide primers:

JAX Universal Mouse Forward: CTAGGCCACAGAATTGAAAGATCT
JAX Universal Mouse Reverse: GTAGGTGGAAATTCTAGCATCATCC
HygR Primer Set 2 Forward: GCTCAGGCACTGGATGAACT
HygR Primer Set 2 Reverse: CAGCCAGTTCTGGGTGTCTT

12.5 μl of PCR reactions were run in agarose gel electrophoresis and stained with ethidium bromide and remaining PCR sample was stocked as preamplified DNA. Samples used in Figures 4D were prepared by reamplification of 1:200 diluted preamplified DNA using target primer sets; 12.5 μl of that secondary reamplified reaction was run in 2.5% agarose gel electrophoresis.

## Quantification and Statistical Analysis

### RT-qPCR Experiments

For Figure 3A and Figure 7A: 3 BCs, 6 iBCs, and 3 mEpiSC colonies were isolated with unique embryo pipettes and washed with CMF-DPBS and carried through standard Ambion Cells-to-CT protocol including optional DNAse I treatment. 50 μl of sample lysate was used for a 125 μl reverse transcription reaction, then diluted to 140 μl with nuclease free water. RT-qPCR was prepared for each sample in duplicate with TaqMan probes using 4 μl of sample cDNA in 20 μl reactions using TaqMan Gene Expression Mastermix. Detection was prepared on StepOne Plus under standard cycling conditions with *Gapdh* samples on each plate. Plate data was analyzed with Applied Biosystems

Expression Suite v1.1. Figure S2C experiments were performed similarly, except using earlier stage iBCs and BCs, and diluting 113 μl of the reverse transcription reaction product with 40 μl of nuclease free water to ensure enough overage for RT-qPCR.

For Figure 6B: iBC/iBC-PC derived outgrowths in 2iLIF media were cultured and passaged as neat colonies. C57BL/6N-CJL mouse ES cells were cultured similarly in 2iLIF media for positive control, and XGFP mEpiSC were cultured in MCM for negative control. Biological replicates of each culture were sourced for total RNA via QIAzol purification methods and finalized in 40 μl of nuclease free water. cDNA was prepared from total RNAs using Superscript Reverse Transcriptase III FS Kit with Random Hexamers protocol. RT-qPCR was carried out in 20 μl reactions with 2 μl of cDNA of each sample in triplicate for all TaqMan probes with standard fast reaction protocols in TaqMan Universal Fast Mastermix on StepOne Plus. Plate data was analyzed with Applied Biosystems Expression Suite v1.1 and expression data was visualized in Microsoft Excel.

For Figure 7C: iBC generation and control cell cultures were washed with PBS then total RNA was prepared with QIAzol standard techniques finalized in Ambion Nuclease Free Water. cDNA was prepared from total RNA using Superscript Reverse Transcriptase III FS Kit with Random Hexamers protocol and diluted with nuclease free water. RT-qPCR was prepared for each sample in triplicate with TaqMan probes for 20 μl reactions using Taqman Gene Expression Mastermix. Detection was prepared on StepOne Plus under standard cycling conditions with *Gapdh* samples as control. Plate data was analyzed with Applied Biosystems Expression Suite v1.1 and expression data was visualized in Microsoft Excel.

For Figure S1C: Naive conversion and control cell cultures were washed with PBS then total RNA was prepared with TRIzol standard techniques and finalized in Ambion RNAsecure. cDNA was prepared from total RNAs using Superscript Reverse Transcriptase III FS Kit with Random Hexamers protocol. RT-qPCR was prepared for each sample in triplicate with standard fast reaction protocols in TaqMan Universal Fast Mastermix for 10ul reactions with TaqMan probes in 384W plate and run on Applied Biosystems 7900HT. Plate data was analyzed with SDS software and expression data was visualized in Microsoft Excel.

